# Sox17 and β-catenin co-occupy Wnt-responsive enhancers to govern the endodermal gene regulatory network

**DOI:** 10.1101/2020.02.19.956565

**Authors:** Shreyasi Mukherjee, Praneet Chaturvedi, Scott A. Rankin, Margaret B. Fish, Marcin Wlizla, Kitt D. Paraiso, Melissa MacDonald, Xiaoting Chen, Matthew T. Weirauch, Ira L. Blitz, Ken W. Y. Cho, Aaron M. Zorn

## Abstract

Lineage specification is governed by gene regulatory networks (GRNs) that integrate the activity of signaling effectors and transcription factors (TFs) on enhancers. Sox17 is a key transcriptional regulator of definitive endoderm development, and yet, its genomic targets remain largely uncharacterized. Here, using genomic approaches and epistasis experiments, we define the Sox17-governed endoderm GRN in *Xenopus* gastrulae. We show that Sox17 functionally interacts with the canonical Wnt pathway to specify and pattern the endoderm while repressing alternative mesectoderm fates. Sox17 and β-catenin co-occupy hundreds of key enhancers. In some cases, Sox17 and β-catenin synergistically activate transcription independent of Tcfs, whereas on other enhancers, Sox17 represses ß-catenin/Tcf-mediated transcription to spatially restrict gene expression domains. Our findings establish Sox17 as a tissue-specific modifier of Wnt responses and point to a novel paradigm where genomic specificity of Wnt/ß-catenin transcription is determined through functional interactions between lineage-specific Sox TFs and ß-catenin/Tcf transcriptional complexes. Given the ubiquitous nature of Sox TFs and Wnt signaling, this mechanism has important implications across a diverse range of developmental and disease contexts.

**Key findings:**

- Sox17 regulates germ layer segregation by promoting endoderm differentiation while simultaneously repressing mesectoderm fates.
- Functional interactions between Sox17 and canonical Wnt-signaling is a major feature of the GRN controlling endoderm specification and patterning.
- Sox17 and β-catenin co-occupy a subset of enhancers to synergistically stimulate transcription in the absence of Tcfs.
- Sox17 regulates the spatial expression domains of Wnt/β-catenin-responsive transcription in the gastrula.

## INTRODUCTION

During embryogenesis, the pluripotent zygote progressively gives rise to specialized cell types expressing distinct sets of genes that, in turn, define the cell’s identity and encode proteins necessary for its function. Lineage specific gene expression is controlled by the genomic integration of signaling pathways and transcription factors (TFs) on DNA cis-regulatory modules (CRMs), such as enhancers, that control transcription (Heinz et al., 2015; Stevens et al., 2017). An important goal of developmental biology is to elucidate how signaling effectors and TFs interact on distinct sets of CRMs within the chromatin landscape to form gene regulatory networks (GRNs) that activate lineage-specific transcriptional programs whilst repressing expression of alternative fates (Charney et al., 2017b). This will provide a deeper understanding of how transcriptional networks are established, dysregulated in disease, and how GRNs might be manipulated for therapeutic purposes.

We have addressed this in the context of endoderm germ layer specification: one of the earliest cell fate decisions in vertebrate development that provides a relatively simple model to elucidate how GRNs control lineage-specific transcriptional programs (Charney et al., 2017b). In embryos and pluripotent stem cells (PSCs), Nodal/Smad2 and Wnt/ß-catenin (Ctnnb1; hereafter Bcat) signaling cooperate to initiate the mesendoderm program and subsequent development of definitive endoderm progenitors, which gives rise to the epithelia of the digestive and respiratory systems (Sumi et al., 2008; Zorn and Wells, 2009). Downstream of Nodal and Wnt, a core set of endoderm TFs: Sox17, Gata4-6, Eomes, Foxa1/2 and Mix family of homeodomain TFs execute the endoderm GRN (Arnold et al., 2008; Charney et al., 2017b; Engert et al., 2013; Sinner et al., 2006). In *Xenopus* where endoderm specification is well studied, the Nodal and Wnt pathways interact at several levels. First, maternal (m) Wnt/Bcat, active on the dorsal side of the blastula, is required for high levels of *nodal* expression at the onset of zygotic transcription (Hyde and Old, 2000). Then Nodal and mWnt cooperate to promote the expression of endoderm TFs Sox17, Foxa2 and many dorsal mesendoderm organizer genes (Xanthos et al., 2002). A few hours later zygotic (z) *wnt8* (a Nodal target) is expressed on the ventral side of the gastrula where, together with Nodal signaling, it promotes ventral and posterior mesendoderm identities (Charney et al., 2017b; Stevens et al., 2017). In mammals early Wnt/Bcat similarly activates expression of Nodal and core endoderm TFs, with prolonged Wnt promoting posterior hindgut fate (Engert et al., 2013; Zorn and Wells, 2009).

Functional interactions between the core endoderm TFs and the Wnt pathway are thought to 1) segregate the transient mesendoderm into endoderm and mesoderm, 2) pattern the nascent endoderm into spatially distinct subtypes and 3) execute the downstream differentiation program to give rise to endoderm-derived lineages. But mechanistically, how the Wnt signaling machinery interacts with core endoderm TFs to execute this differentiation cascade is unresolved. The transcriptional targets of the endoderm TFs are largely unknown, and it is unclear exactly how Wnt/Bcat regulates distinct spatiotemporal transcription programs in the embryo. Indeed, how the canonical Wnt pathway elicits context-specific transcriptional responses in its multitude of different biological roles from development, homeostasis and cancer is still poorly understood. According to the dogma of canonical Wnt signaling, Wnt-activated Fzd-Lrp5/6 receptor complexes sequester the Bcat degradation complex, resulting in the stabilization and translocation of Bcat to the nucleus. There, it interacts with one of four Tcf HMG-box TFs (Tcf7, Tcf7L1, Tcf7L2 and Lef1) at Wnt-responsive CRMs (Cadigan and Waterman, 2012; Schuijers et al., 2014), ultimately activating a multiprotein “Wnt-enhanceosome” with the scaffold proteins Bcl9, Pygopus and the ChiLS complex (Gammons and Bienz, 2018). In the absence of Bcat, Tcfs are coupled with corepressor proteins Tle/Groucho and histone deacetylase Hdac to inhibit transcription of Wnt target genes. Bcat displaces Tle and recruits a co-activator complex including histone acetyltransferases Ep300 or Cbp to stimulate transcription (Cadigan and Waterman, 2012). How distinct context-specific Wnt target genes are selected is unclear since all Tcfs have nearly identical DNA-binding specificities (Badis et al., 2009; Ramakrishnan and Cadigan, 2017) and for the most part they are ubiquitously expressed. An emerging idea is that the Wnt-enhanceosome also interacts with other lineage-specific TFs to integrate lineage-specific inputs (Gammons and Bienz, 2018; Trompouki et al., 2011); yet, how these impact genomic specificity *in vivo* is largely untested and the idea of Tcf-independent Wnt-mediated transcription remains controversial.

In this study we investigated the possibility that Sox17 functionally interacts with Wnt/Bcat to regulate transcription in the *Xenopus* endodermal GRN. In all vertebrate embryos, Sox17 is specifically expressed in the gastrula endoderm where it is required for early gut development (Clements et al., 2003; Hudson et al., 1997; Kanai-Azuma et al., 2002; Viotti et al., 2014). Despite the critical role of Sox17 in endoderm development, only few of its direct transcriptional target genes have been identified (e.g.: *hnf1b, foxa1* and *dhh*) (Ahmed et al., 2004; Sinner et al., 2004; Yagi et al., 2008). In *Xenopus*, ectopic Sox17 is sufficient to initiate endoderm development in pluripotent blastula animal cap cells (Clements et al., 2003; Hudson et al., 1997), and co-injection of stabilized Bcat can enhance this activity (Sinner et al., 2004). Sox17 can physically interact with Bcat *in vitro* and suppress the transcriptional activity of generic Tcf/Bcat reporter constructs (TOPflash) in tissue culture experiments (Sinner et al., 2007; Zorn et al., 1999). However, the biological relevance of these interactions and whether Sox17 and Bcat functionally interact on chromatin to regulate the endoderm GRN remains unknown.

Here, we defined the genomic targets of Sox17 in the *Xenopus* gastrula. In addition to promoting expression of endoderm genes, Sox17 also represses ectoderm and mesoderm gene transcription, and acts in a negative feedback loop to restrain Nodal signaling. We demonstrate that functional interaction with canonical Wnt signaling is a key feature of the Sox17-regulated GRN. Over a third of all Bcat and Sox17 genomic binding in the gastrula occur at the same CRMs. In some instances, Sox17 suppressed Bcat-Tcf mediated transcription, while in other cases, Sox17 and Bcat synergistically activated enhancers apparently independently of Tcfs, indicating a novel mode of regulation. These results provide new insight into the GRN controlling endoderm development and have implications for how Sox TFs and Bcat might interact in diverse biological contexts from development to cancer.

## RESULTS

### Sox17 regulates a genomic program controlling germ layer segregation and endoderm development

To identify the transcriptional program regulated by Sox17, we performed RNA-sequencing (RNA-seq) on control and Sox17-depleted *Xenopus tropicalis* embryos at multiple time points during blastula and gastrula stages (NF9-12) when the endoderm germ layer is being specified. In *Xenopus tropicalis* there are three redundant genes: *sox17a, sox17b. 1* and *sox17b.2* (collectively *sox17*) with indistinguishable activities and identical expression in presumptive vegetal endoderm cells of gastrula embryos (Fig. 1A and Fig. S1A) (D’Souza et al., 2003; Hellsten et al., 2010). Microinjection of a combination of antisense morpholino oligos (sox17aMO and sox17bMO) targeting all 3 paralogs (Fig. S1B) resulted in a robust knockdown of Sox17 protein as confirmed by immunostaining (Fig. 1C,B). The Sox17-MO phenotype was consistent with previous reports (Clements et al., 2003; Sinner et al., 2006) and phenocopied mouse mutants with defective gut development (Kanai-Azuma et al., 2002). Injection of mRNA encoding mouse Sox17 rescued both the anatomical and gene expression phenotypes confirming the efficacy and specificity of the MOs (Fig. S1C,D).

**Fig. 1.**
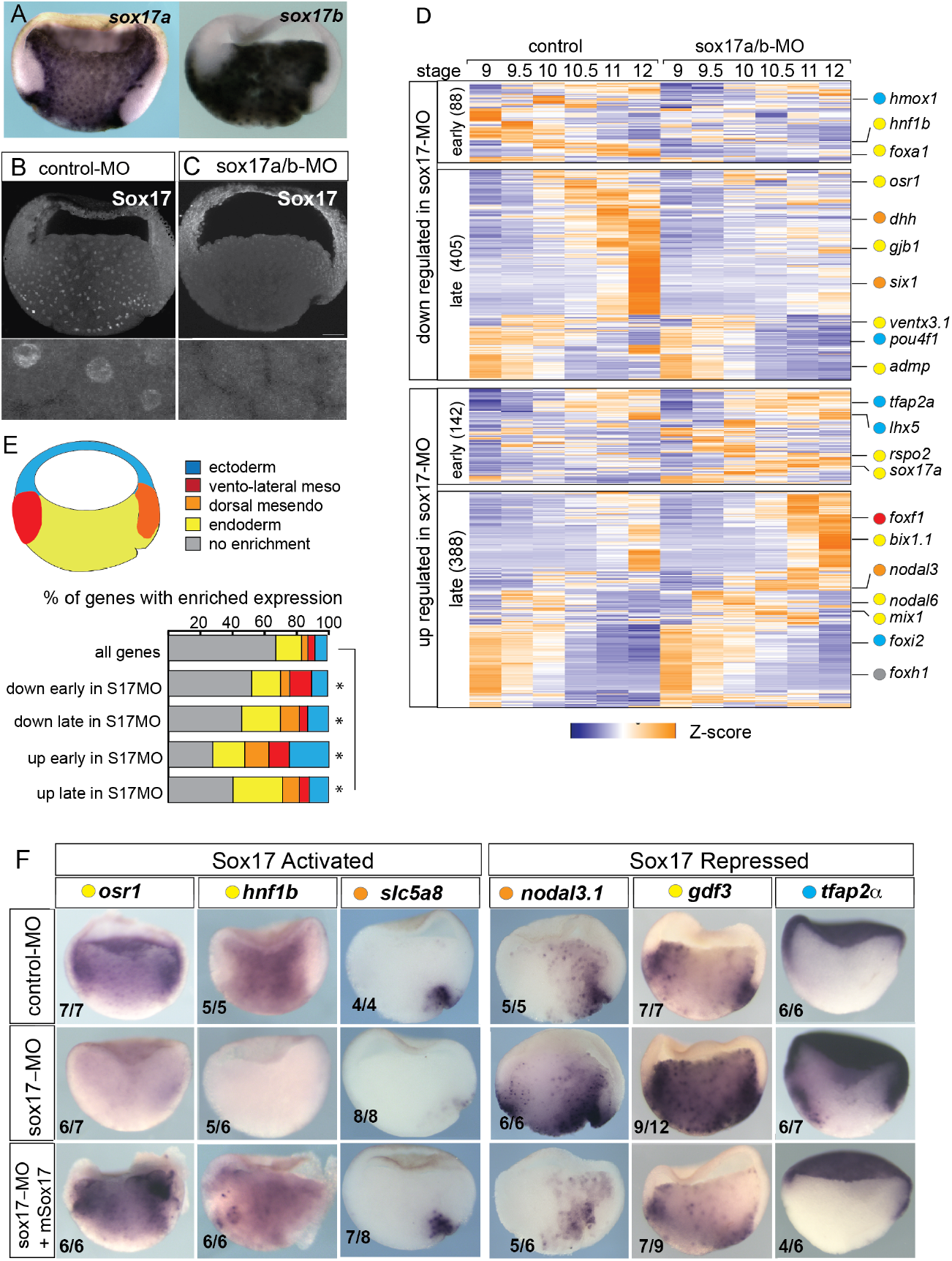
Sox17-regulated endoderm transcriptome. **(A)** Endoderm expression of *sox17a* and *sox17b* in NF10.5 *Xenopus tropicalis* gastrula. **(B-C)** Sox17 immunostaining in control-MO (B) and sox17a/b-MO (C) injected gastrula with an antibody that recognizes both Sox17a and Sox17b shows MO effective knockdown. **(D)** Time course heatmap of Sox17-regulated transcripts. Differential expressed genes from a pairwise comparison of control-MO and sox17-MO (≥2 fold change, FDR≤5%) at each stage showing transcripts that are down regulated or up regulated early (NF9-10) or late (NF10.5-12). Key genes are color coded based on regional expression from fate map in (E). **(E)** Enriched expression of Sox17-regulated transcripts. Gastrula fate map (dorsal right) colored by different tissues. Histogram showing enriched expression in differnet regions of the embryos based on RNA-seq of dissected gastrula tissues (Blitz et al., 2017). *p<0.05 Chi squared test relative to all genes in the genome. **(F)** In situ validation of Sox17-regulated transcripts. Disrupted expression in sox17-MOs is rescued by co-injeciton of mouse (m) *Sox17* RNA.

Differential expression analysis of control-MO and Sox17-MO embryos identified 1023 Sox17-regulated genes (≥ 2-fold change, FDR≤ 5%) (Fig. 1D and Table S1). Gene Ontology (GO) enrichment was consistent with the Sox17-regulated transcriptome being involved in “endoderm formation”, “epithelial differentiation” and “digestive track morphogenesis” (Fig. S1E). In total 493 genes were downregulated in Sox17-depleted embryos and 530 genes were upregulated. The time course data revealed that >75% of the differentially expressed genes were Sox17-dependent during the mid-late gastrula (NF10.5-12), consistent with a role in maintaining the endoderm fate after initial induction by Nodal signaling in the blastula (NF9) (Fig.1D and Fig. S1A). Sox17-regulated genes include 73 TFs, including known targets *foxa2* and *hnf1b* (Sinner et al 2004; Sinner et al 2006), and paracrine signaling components (Table S1) such as the Hedgehog pathway (*dhh*, *hhip, gli1*); a key epithelial signal in gut organogenesis. This confirms that Sox17 sits atop of the regulatory hierarchy regulating endoderm differentiation.

We next investigated how Sox17-regulated genes were spatially expressed, leveraging a previously published RNA-seq of different tissues dissected from gastrula embryos (Blitz et al., 2017). As predicted, a majority of Sox17-dependent genes were enriched in the vegetal endoderm and dorsal mesendoderm (organizer) (Fig. 1E) including *osr1, hnf1b, dhh* and *slc5a8* (Fig. 1F and Fig. S1D). Interestingly, over 30% of the genes upregulated early (NF9-10) in the Sox17-depleted embryos were normally enriched in the ectoderm or mesoderm tissue; examples include ectoderm-promoting TFs *tfap2a* and *lhx5* (Houston and Wylie, 2003) (Fig. 1D-F) as well as the mesoderm TFs *foxf1, tbx20* and *hlx*. This suggests that Sox17 plays an important role in repressing ectoderm and mesoderm fate in vegetal endoderm cells. Unexpectedly Sox17 also negatively regulated ~150 genes that are normally enriched in the vegetal endoderm and dorsal mesendoderm. Some of these vegetally enriched genes encoded components of the endoderm-promoting Nodal pathway including *nodal1, nodal6, gdf3, gdf6, foxh1, mix1* and *bix1.1*, all of which were upregulated in Sox17-depleted embryos (Fig. 1D-F and Fig. S1A,D). Indeed, even *sox17* transcripts were modestly increased in Sox17-MO embryos even though previous work has demonstrated that Sox17 directly maintains its own transcription (Howard et al., 2007) after initial induction by Nodal/Smad2. These data indicate that Sox17 acts as a negative feedback inhibitor that restricts excessive endoderm development by restraining Nodal activity after initial induction.

The observation that over 10% of Sox17-regulated genes are enriched in the organizer mesendoderm while Sox17 is present throughout the endoderm, suggests that Sox17 might also functionally interact with the Wnt and/or Bmp dorso-ventral patterning pathways to control spatial expression. Intriguingly, we found that Sox17 regulates the expression of several Wnt/Bcat pathway components and targets including: *dkk1, dkk2, fzd5, nodal3.1, pygo1, sia1, ror1, rsop2* and *wnt11* (Fig. 1D,F and Table S1). Together these data suggest that Sox17 regulates both endoderm specification and patterning while suppressing mesectoderm fate and that Sox17 participates in feedback loops with the Wnt and Nodal pathways.

### Sox17 ChIP-Seq reveals direct endodermal targets and a Nodal and Wnt feedback

To identify direct Sox17-targets we generated and validated anti-Sox17 antibodies (Fig. S2) and performed ChIP-seq of gastrula (NF10.5) embryos, identifying 8436 statistically significant Sox17-bound CRMs (IDR; p<0.05). These were associated with 4801 genes (Fig. 2A), based on annotation to the nearest transcription start-site (TSS) by HOMER. 88% of Sox17-bound loci were in introns or intergenic regions more than 1kb away from the TSS, consistent with distal CRMs (Fig. S3A). A comparison to published ChIP-seq data of *Xenopus* embryos at the same stage (Hontelez et al., 2015) showed that most of the Sox17-bound genomic loci were also bound by Ep300, (Fig. 2B, E) indicative of active enhancers. Motif analysis of the ChIP-seq peaks confirmed that Sox17 motifs were the most enriched, as expected (Fig. 2C and Fig. S3B).

**Fig. 2.**
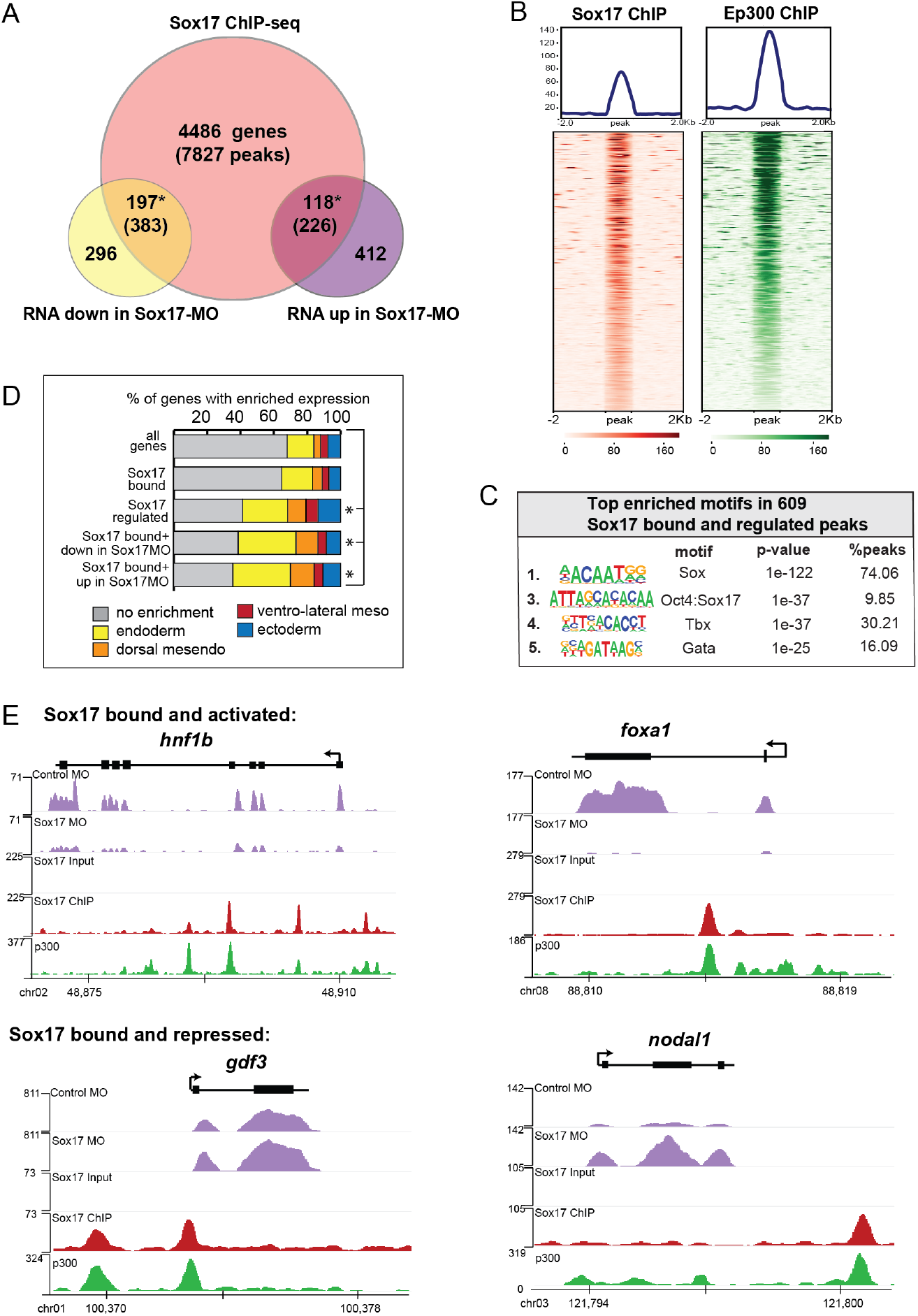
ChIP-seq identifies Sox17-bound enhancers and direct transcriptional targets. Sox17 ChIP-seq of gasturla embryos. Venn diagram of Sox17-bound genes (and associated peaks) from ChIP-seq intersected with Sox17-regulated genes. *p<0.001, hypergeometric test. **(B)** Peak density plots showing that the Sox17-bound loci are cobound by the histone acetyletransferase Ep300, a marker of active enhancers. Average density of all peaks in top panel. **(C)** Motif analysis of TF DNA-binding sites enriched in Sox17 peaks. **(D)** Sox17-bound genes are enriched in the endoderm. The Histogram shows the regional expression pattern of Sox17-bound and regulated genes based on RNA-seq of dissected gastrula tissues (Blitz et al., 2017). p<0.05 Chi squared test relative to all genes in the genome. **(D)** Genome browser views of representative genes showing RNA-seq expression in control-MO and sox17-MO embryos and the ChIP-seq tracks of Sox17 and Ep300 bound genomic loci.

A comparison to published human ChIP-Seq data revealed that 20% of the *Xenopus* Sox17-bound genes were also SOX17-bound in PSC-induced definitive endoderm (Fig. S3) (Tsankov et al., 2015). GO analysis of the conserved SOX17-bound genes show an enrichment for ‘Tgfb receptor activity’ and ‘Bcat binding’ (Fig. S3) reinforcing the notion that functional interaction with the Nodal and Wnt pathways is a conserved feature of the Sox17-regulated endoderm GRN.

Intersecting the RNA-seq and ChIP-seq data identified 315 genes associated with 609 Sox17-bound enhancers, which are likely to be direct transcriptional targets (Fig. 2A). These putative direct targets had significantly enriched (44%, 139/315 transcripts) expression in the endoderm or dorsal mesendoderm (Fig. 2D). Of the Sox17-bound and regulated genes, 197 genes (383 peaks) were positively regulated by Sox17 (down in Sox17MOs) including *slc5a8, hnf1b, foxa1* and *dhh* (Fig. 2E), all of which were previously suggested to be direct Sox17-targets. Sox17 negatively regulated 118 putative direct targets (up in Sox17MOs) including ectodermal genes (*lhx5, foxi2* and *tfap2a*), endoderm-enriched Nodal pathway genes (*nodal1, gdf3, gdf6* and *mix1*) and Wnt-regulated organizer genes (*dkk* and *fst*), suggesting direct transcriptional repression (Fig. 2E). Interestingly ~45% of peaks associated with Sox17-activated genes were also enriched for LIM-homeodomain binding sites, in contrast to peaks from Sox17-repressed genes which were enriched for Tbx or Pou motifs (Fig. S3B). This suggests that Sox17 may coordinately engage enhancers with other core endoderm GRN TFs and mediate activation or repression of target genes depending on the interacting TFs.

These analyses provide new insight into the endoderm GRN and reveal previously unappreciated roles for Sox17 in germ layer segregation and endoderm patterning involving functional interactions with the Nodal and Wnt pathways. These findings, together with our previous work demonstrating that Sox17-Bcat can physically interact *in vitro* (Sinner et al., 2007; Zorn et al., 1999) prompted us to investigate their genomic interactions.

### β-catenin directly regulates the endodermal transcriptome

To test the hypothesis that Sox17 and Bcat functionally interact *in vivo*, we set out to identify Bcat-regulated genes and compare these to the Sox17-regulated transcriptome. Injection of a well-characterized Bcat-MO (Heasman et al., 2000) resulted in depletion of nuclear Bcat, reduced expression of a transgenic Wnt-responsive reporter (Tran et al., 2010) and the expected ventralized phenotype, which was rescued by co-injection of stabilized human Bcat mRNA (Fig. S4), thus confirming the efficacy and specificity of the Bcat knockdown. RNA-seq of control-MO and Bcat-MO depleted embryos at eight time points from blastula and gastrula stages (NF7-12) identified a total of 2568 Bcat-regulated genes (≥ 2-fold change, FDR ≤ 5%), 1321 of which were downregulated and 1247 upregulated in the Bcat-MO embryos (Fig. 3A and Table S2). Remarkably the Bcat-dependent genes encoded 251 TFs, (~20% of all the TFs in the genome), reinforcing the notion that Wnt/Bcat initiates a transcriptional cascade in the early embryo.

**Fig. 3.**
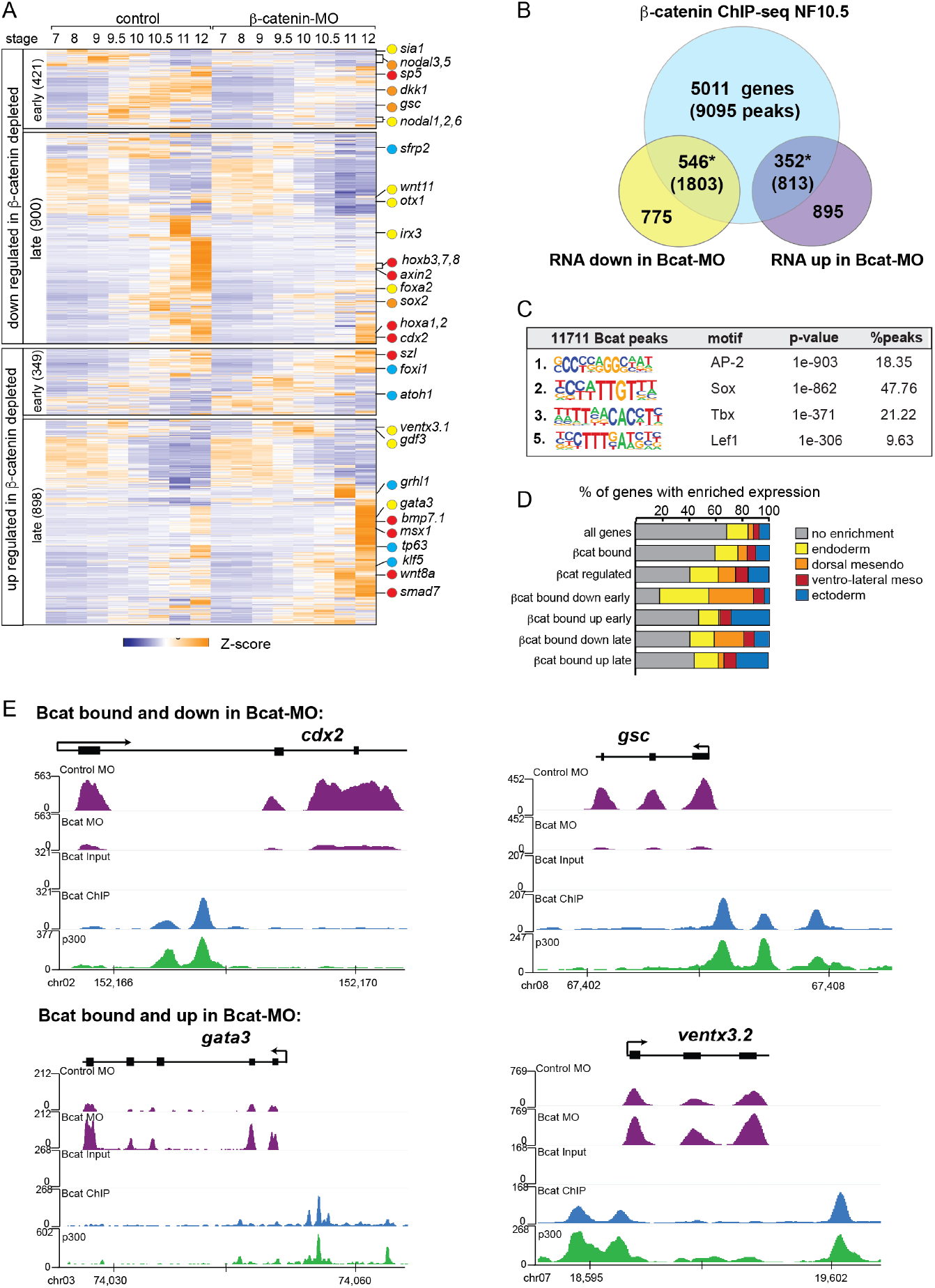
The Bcat regulated genomic program. **(A)** Time course heatmap of Bcat-regulated transcripts. Differential expressed genes from a pairwise comparison of control-MO and Bcat-MO embryos (≥2 FC, FDR≤5%) at each stage identified 2568 Bcat-regulated transcripts; 1321 downregulated and 1247 upregulated in early (NF7-9.5) or late (NF10-12) Bcat-MO embryos. Key genes are indicated color coded based on regional expression from (D). **(B)** Venn diagram of Bcat-bound genes associated with Bcat peaks from published ChIP-seq (Nakamura et al., 2016) intersected with Bcat-regulated genes identified 898 putative direct targets associated with 2616 peaks. *p<0.001, hypergeometric test. **(C)** Motif enrichment analysis of Bcat peaks. **(D)** Histogram showing the expression pattern of Bcat-bound and regulated genes based on RNA-seq of dissected gastrula tissues (Blitz et al., 2017). **(E)** Genome browser views of representative genes showing the RNA-seq expression in control-MO and Bcat-MO embryos and the ChIP-seq tracks of Bcat and Ep300 bound genomic loci.

Intersecting the Bcat-MO RNA-seq data with previously published Bcat ChIP-seq data from *Xenopus tropicalis* gastrula (Nakamura et al., 2016) identified 898 putative direct target genes associated with 2616 Bcat bound CRMs. 546 genes were downregulated in the Bcat-MO and 352 genes had increased expression in Bcat-MO embryos (Fig. 2B). These included almost all of the previously known Bcat targets in early *Xenopus* embryos and had extensive overlap with other recent genomic analysis of Wnt targets in the *Xenopus* gastrula (Ding et al., 2017; Kjolby and Harland, 2017; Nakamura et al., 2016) (Fig. S5B).

As expected, many of the positively regulated Bcat targets had enriched expression in the dorsal mesoderm (Fig. 3D) including most known organizer genes such as *cer1, chrd, dkk1, frzb, fst and gsc*. In contrast ~30% of Bcat-bound genes with increased expression in the early Bcat-MO embryos (NF7-9) were enriched for ectoderm specific transcripts, consistent with the expansion of ectoderm and loss of neural tissue in ventralized embryos (Fig. 3D). We also identify many direct Bcat targets that are likely to be regulated by zygotic Wnt8, (Kjolby and Harland, 2017; Nakamura et al., 2016) including *sp5, axin2, cdx1/2, fzd10, msgrn1* and eight *hox* genes, all of which were downregulated in Bcat-MOs (Fig. 3A, E). A number of zygotic Wnt-targets with enriched expression in the ventral mesoderm were components of BMP pathway including; *bambi*, *bmp7*, *id2*, *msx1, smad7, szl*, *ventx1*, and *ventx3* (collectively known as the BMP-synexpression group) (von Bubnoff et al., 2005). Most of these genes are known to be directly activated by both Bmp4/Smad1 and zWnt8/Bcat (Itasaki and Hoppler, 2010; Stevens et al., 2017) but were upregulated, rather than downregulated, in Bcat-depleted embryos, likely due to the increase in BMP/zWnt8 signaling in ventralized embryos.

Most importantly, almost 200 Bcat-bound and regulated genes had endoderm enriched expression including most of the core endoderm TFs; *sox17, foxa, eomes, gata4, mix1* and *mixer* (Fig. 3D and Table S2). Thus, contrary to the prevailing view that Bcat promotes endoderm fate primarily by activating Nodal ligand expression (Blythe et al., 2010; Hyde and Old, 2000), we find that Wnt/Bcat also directly regulates the transcription of many endodermal genes. Surprisingly, *de novo* motif analysis showed that the Bcat bound peaks were more enriched for Sox than canonical Tcf motifs (Fig. 3C and Fig. S5A), supporting the hypothesis that Sox17 and Bcat coregulate endodermal transcription.

### Sox17 and β-catenin co-occupy endoderm enhancers

Intersection of the ChIP-seq datasets identified 3956 genomic loci co-occupied by both Sox17 and Bcat, and these had epigenetic signatures of active enhancers (Fig. 4A,B). This represents a third of all Sox17 genomic binding in the gastrula genome. Comparison of the Sox17-MO and Bcat-MO RNA-seq datasets identified 415 transcripts that were regulated by both Sox17 and Bcat; comprising of ~40% of the Sox17-regulated and ~15% of the Bcat-regulated transcriptome (Fig. 4A). These data suggest that co-regulation with Wnt/Bcat is a major feature of the endoderm GRN.

**Fig. 4.**
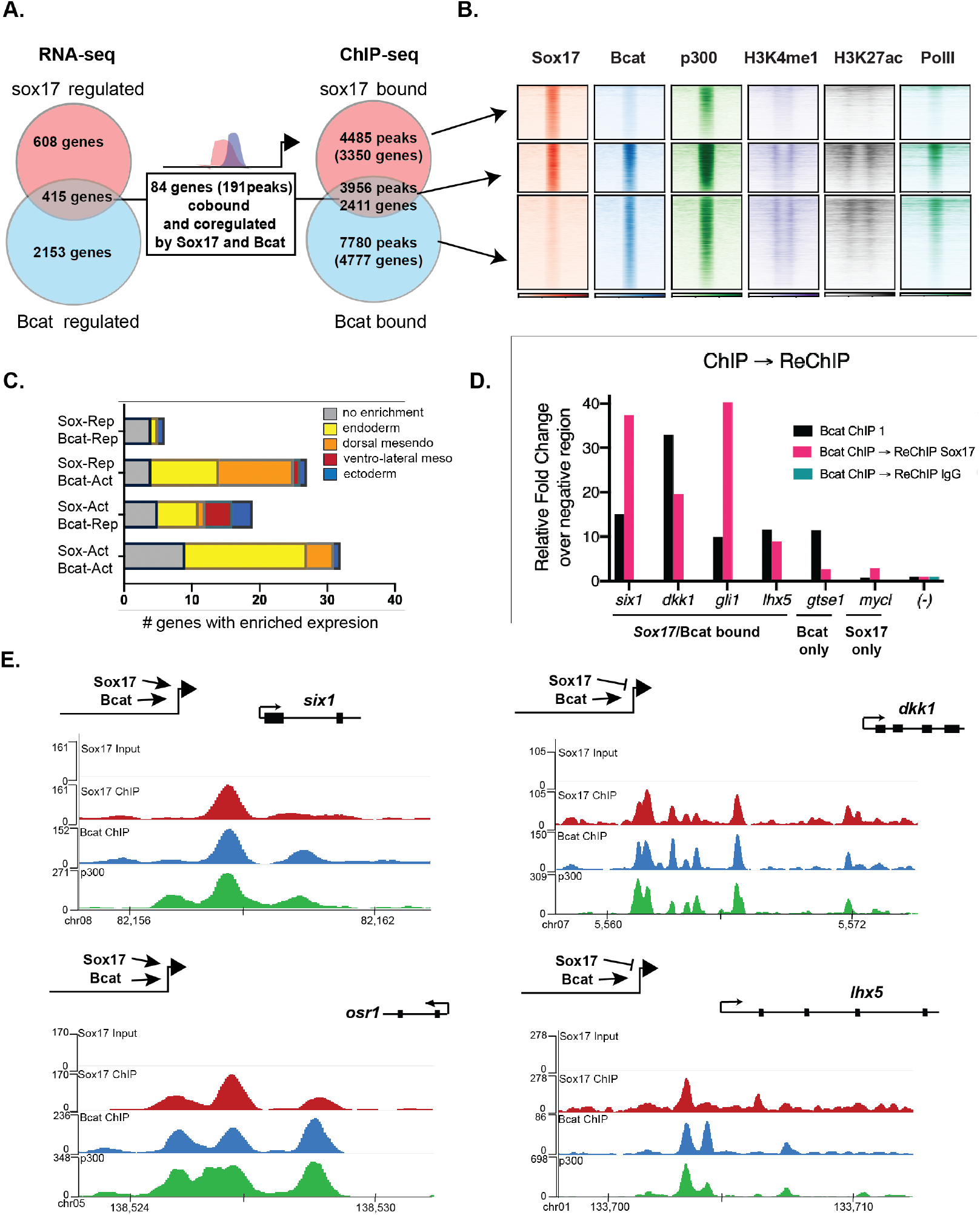
Sox17 and Bcat co-bind enhancers to regulated the endoderm GRN. **(A)** 84 genes (191 Peaks) are cobound and coregulated by Sox17 and Bcat. Venn diagram shows that 416 genes are regulated by both Sox17 and Bcat (left), whereas 2411 genes (3956 peaks) are bound by both Sox17 and Bcat. **(B)** Density plots showing that Sox17 and Bcat cobound peaks are enriched for active enhancers marker; Ep300, H3K4me1 and H3K27Ac and RNA pol II. **(C)** Regional enrichment of all subsets of enhancers cobound and coregulated by Sox17 and Bcat. **(D)** ChIP-reChIP experiments show that Sox17 and Bcat coocupy all the test CRMs (*six1, dkk1, gli1* and *lhx5*) in the same cells, in contrast to the *gtse1* loci, which is bound only by Bcat. **(E)** Genome browser views of representative genes and the ChIP-seq tracks showing the genomic loci bound by Bcat and Ep300.

In total, 84 genes (191 Peaks) were bound and regulated by both Sox17 and Bcat (Fig. 4A,E). This is likely to be an underestimation of the number of direct coregulated genes, as these had to pass a stringent thresholding of four independent statistical tests to be included in this list. A comprehensive motif analysis of the 191 co-bound and coregulated peaks using the CIS-BP database (Lambert et al., 2019) revealed that 78% (149/191) of the peaks contained both Sox17 and Tcf DNA-binding motifs, often > 5 sites in each peak (e.g., *dkk1, lhx5* and *osr1*) (Fig. S6 B,D, Table S5). In contrast 21% (40/191) of the peaks had Sox motifs but no obvious Tcf consensus sites (e.g., *six1*), suggesting that Bcat might be recruited to these loci independent of Tcfs. Only 1/191 peaks had Tcf but no Sox motifs. In most cases individual genes were associated with multiple Sox17 and Bcat co-occupied peaks, suggesting that combinatorial binding and integration of several CRMs is crucial to control transcription (Fig. 4E).

Since the ChIP-seq experiments were performed on whole embryos, it was possible that Sox17 and Bcat chromatin binding might occur in different cell populations. To test whether Sox17 and Bcat bound the same enhancers in the same cells we performed ChIP-reChIP experiments. First anti-Bcat ChIP was performed on 500 gastrulae. DNA eluted from the first ChIP was then re-precipitated with either anti-Sox17 or a negative control IgG antibody. All 4 loci tested by ChIP-reChIP QPCR (*six1, dkk1, lhx5* and *gli1*) showed enrichment compared to IgG demonstrating that these genomic loci (~150 bp in length) were simultaneously bound by Sox17 and Bcat in endodermal cells. In contrast a negative control locus *gtse1* which is bound by Bcat but not Sox17 was not recovered in the ChIP-reChIP experiment (Fig. 4D).

The 84 co-bound and coregulated genes fell into 4 regulatory categories (Fig. 4C, E); 32/84 were activated by both Sox17 and Bcat (e.g., *six1* and *osr1*) and tended to have endoderm enriched expression, 27/84 were activated by Bcat but repressed by Sox17 (e.g., *dkk1*) and these tended to be enriched in the organizer mesendoderm, 19/84 were positively regulated by Sox17 and negatively regulated by Bcat (up in the Bcat MO) and finally 6/84 appeared to be negatively regulated by both Sox17 and Bcat. The observation that Bcat positively regulated most of these genes (59/84, 70%) was consistent with its known role in transcriptional activation. Sox17 positively regulated 60% of targets and repressed 40% of targets, suggesting context-dependent activity. For further analysis we focused on the two major regulatory groups; genes/enhancers activated by both Sox17-Bcat and activated by Bcat but repressed by Sox17 (Fig. 4E).

### Sox17 and β-catenin coordinate spatial expression domains in the embryo

It was possible that the overlap in Sox17- and Bcat-regulated genes was largely due to the Sox17-Wnt feedback loop where Bcat is required for robust *sox17* expression and Sox17, in turn, regulates the expression of Wnt-pathway components. While this is likely to be part of the mechanism, we directly tested the functional necessity for both Sox17 and Bcat in epistasis experiments, asking whether injection of *sox17* RNA could rescue coregulated genes in Bcat depleted embryos and vice versa. Co-injection of *mSox17* mRNA could not rescue the normal expression patterns of *six1, osr1, lhx5* or *dkk1* in Bcat-MO embryos, but it did rescue *foxa1* expression (Fig. 5). This demonstrates that reduced Sox17 in Bcat depleted embryos cannot account for the disrupted *six1, osr1, lhx5* or *dkk1* expression, but it can account for the reduced *foxa1* expression. Similarly, co-injection of stabilized (*ca*)*Bcat* mRNA rescued the normal expression of *foxa1* but not *six1, osr1, lhx5* or *dkk1* in Sox17-morphants indicating that the defects in their expression were not simply due to reduced Wnt signaling (Fig. 5).

**Fig. 5.**
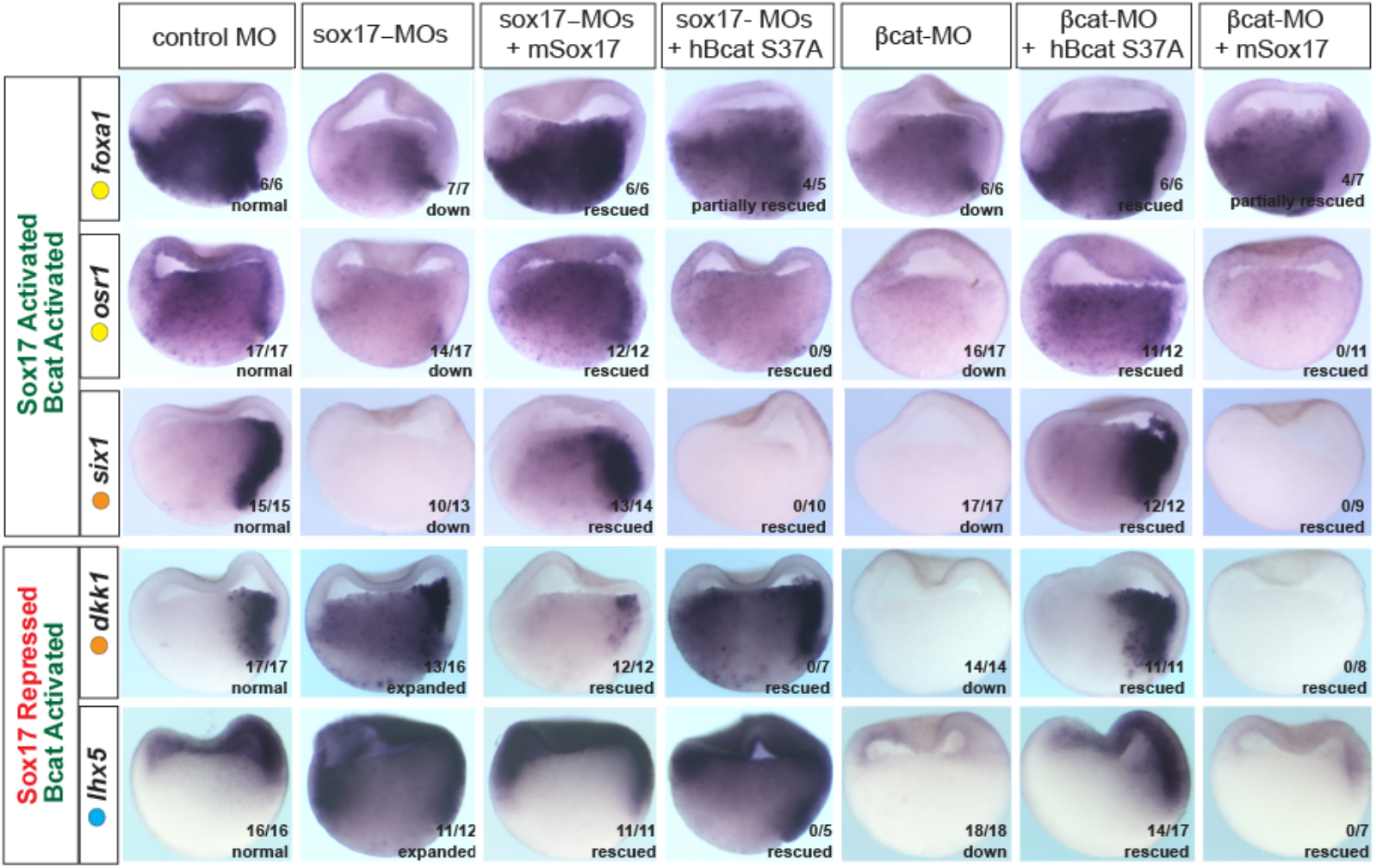
Sox17 and Bcat epistatsis. In situ hybridization of gastrula embryos (dorsal right) injected with sox17-MO or bcat-MO some of which were co-injected with RNA encoding either mSox17 or a human S37A stabilized Bcat. The genes *osr1, six1* and *foxa1* are activated by both Sox17 and Bcat, whereas *dkk1* and *lhx5* are activated by Bcat but repressed by Sox17. Co-injection of mSox17 cannot resuce normal expression in Bcat-MOs, nor can hBcat S37A RNA rescue Sox17-MOs indicating that both Sox17 and Bcat required for normal expression.

These experiments also show that Sox17 and Bcat coordinate spatial gene expression in the embryo in a complex gene-specific manner. Wnt/Bcat stimulates expression of *lhx5* in the ectoderm, but Sox17 represses this activity in the presumptive endoderm. Sox17 and Bcat are both required for robust expression of *osr1* and *six1* in the endoderm and dorsal mesendoderm respectively. In contrast Sox17 normally suppresses Bcat-activated expression of *dkk1* in the deep endoderm. Together, with their co-occupancy at CRMs, these data show that Sox17 and Bcat are both required to directly regulate transcription in discrete spatial domains.

### Sox17 and Bcat synergistically regulate transcription of endodermal enhancers

To examine how Sox17 and Bcat directly regulate endodermal transcription, we selected four exemplar co-occupied enhancers: a) a proximal *six1* enhancer located −1kb from the TSS, b) a proximal *dkk1* enhancer located −1.6kb from its TSS (Fig. 6B), c) a distal −7.5kb *dkk1* enhancer and d) a −1kb *lhx5* enhancer (Fig. S7C, E). We focused most of our analysis on *six1* and *dkk1* which are both expressed in the dorsal mesendoderm but have opposite regulation. Sox17-Bcat promotes *six1*, whilst Sox17 represses Bcat activation of *dkk1* (Fig. 5).

**Fig. 6.**
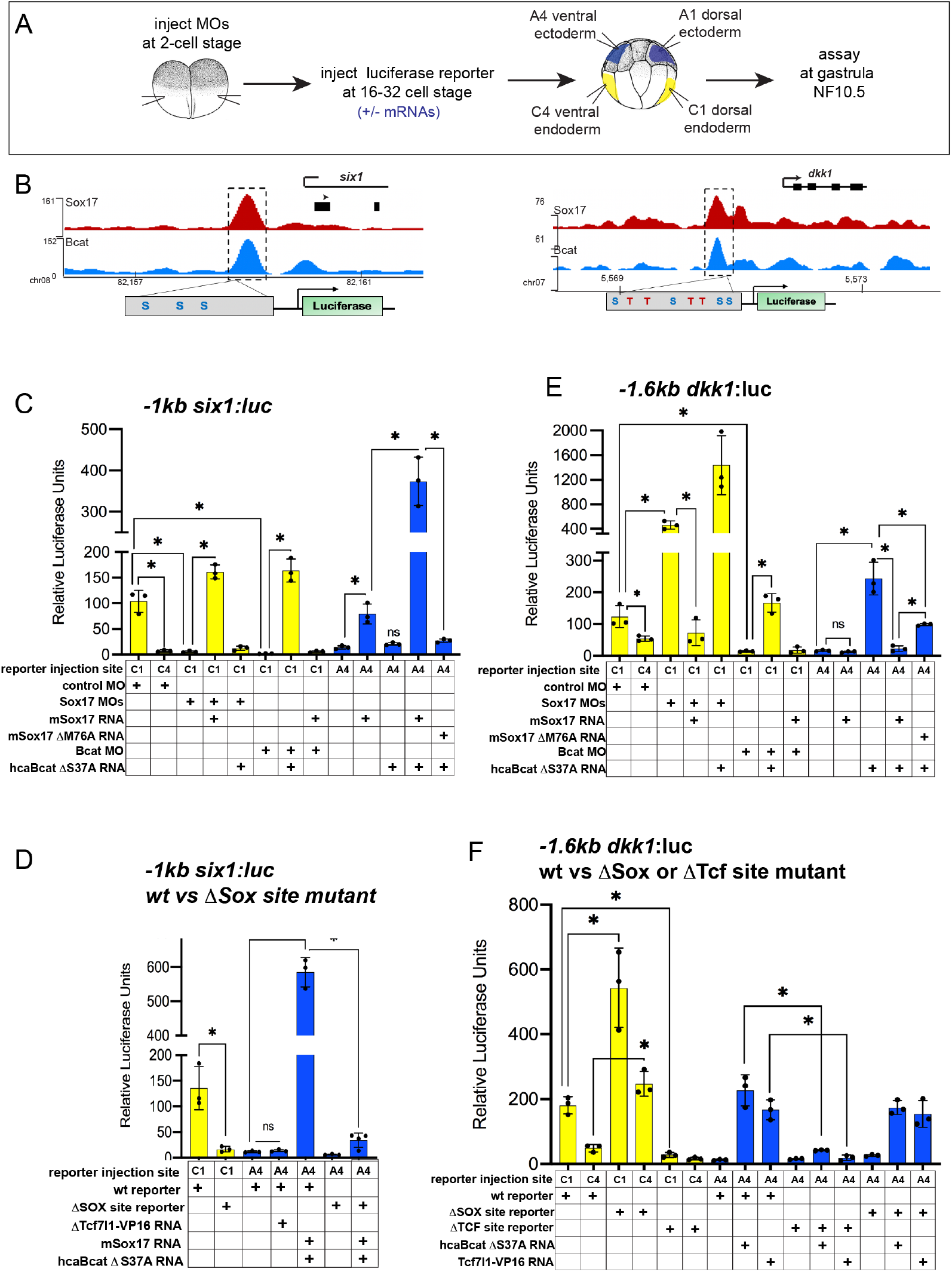
Sox17 and Bcat coordinately regulate enhancers. **(A)** Experimental design. **(B)** IGV genome browser tracks showing Sox17 and Bcat binding and the predicted Sox17 and Tcf binding sites at the −1kb *six1* and-1.6kb *dkk1* enhancers **(C-F)** Luciferase assay of (C) wildtype (wt) −1kb *six1:luc*, (D) wt versus ΔSox-site mutant 1kb *six1:luc*, (E) wt −1.6kb *dkk1:luc* and (F) wt versus ΔSox-site and ΔTcf-site mutant −1.6kb *dkk1:luc* reporters injected in different tissue with the indicated MOs and/or mRNAs. Standard Deivation, *p<0.05 in pairwise Student T-tests.

The −1kb *six1* enhancer is evolutionarily conserved from *Xenopus* to human containing three Sox-binding sites but no predicted Tcf binding sites (Fig. 6B). In mouse, this enhancer can drive transgenic expression in the embryonic gut tube, although the TFs that modulate it were previously unknown (Sato et al., 2012). Analysis of published ChIP-seq data indicate that, like in *Xenopus*, the human *SIX1* CRM is also bound by SOX17 in human PSC-derived endoderm (Fig. S7A) (Tsankov et al., 2015). Mammalian *Dkk1* is also known to be a direct Bcat-Tcf target (Niida et al., 2004), but the enhancers that regulate its expression in the gastrula had been previously uncharacterized. Unlike the −1kb *six1* enhancer, the −1.6kb and −7.5 *dkk1* enhancers as well as the −1kb *lhx5* enhancer harbors both Tcf and Sox DNA-binding sites (Fig. 6B, S7 and file S1).

We cloned each of these CRMs into luciferase (luc) reporter constructs and tested whether they require Sox17 and Bcat for enhancer activity in *Xenopus* embryos (Fig. 6A). Targeted microinjection into different cells of 32-cell-stage *Xenopus* embryos showed that all the reporters recapitulated the endogenous spatial expression indicating that they are *bona fide* enhancers. The *six1* and *dkk1* reporters were both active in the dorsal-anterior endoderm, but not the ventral endoderm or ectoderm (Fig. 6C,E, S7F), whereas the *lhx5* reporter was only active in the dorsal ectoderm (Fig. S7D). Depletion of endogenous Sox17 or Bcat abolished expression of the - *1six1*:luc (Fig.6C), and this was rescued by adding back *mSox17* to Sox17-MO or by adding back *caBcat* mRNA to Bcat-MO embryos. However, co-injection of *mSox17* could not rescue expression in Bcat-MO embryos and nor could *caBcat* RNA rescue expression in Sox17-MOs (Fig. 5, 6C). These epistasis experiments indicate that both Sox17 and Bcat are required for activation of the −1kb *six1* enhancer. The human *SIX1:luc* construct exhibited identical activity to the *Xenopus six1:luc* enhancer (Fig. S7B). In contrast, in the case of the *-1.6kb dkk1*, −7.*5kb dkk1* and the *-1kb lhx5:luc* constructs, depletion of Bcat abrogated reporter activity, whereas Sox17 depletion resulted in significantly elevated reporter activity, indicating that Sox17 restricts Bcat-mediated transactivation of the *dkk1* and *lhx5* enhancers (Fig. 6E, S7D,F).

To test whether Sox17 and Bcat were sufficient to regulate the enhancers, we assayed their activity in ventral ectoderm tissue which does not express endogenous Sox17 and has low Wnt/Bcat activity. Injection of *mSox17* but not *caBcat* mRNA was sufficient to stimulate the *-1kb six1:luc* and human *SIX1:luc* reporters in the ectoderm, but co-injection of *caBcat* along with *mSox17* synergistically activate the reporters (Fig. 6C, S7B). On the other hand, *caBcat* but not *mSox17* was able to activate the *dkk1* and *lhx5* reporters in ventral ectoderm, and co-injection of *mSox17* suppressed the ability of *caBcat* to stimulate these reporters (Fig. 6E, S7D,F), consistent with the endogenous regulation (Fig. 5). The ability of Sox17 to activate the *six1* reporters or suppress Bcat-induced activation of the *dkk1* and *lhx5* reporters required the DNA-binding function of Sox17. A M76A point mutation in mSox17, known to disrupt the DNA-binding HMG domain (Sinner et al., 2007) significantly reduced Sox17’s ability to (Fig. 6C, E, S7B,D,F).

The *six1* enhancer is predicted to have only Sox17 sites with no evidence of Tcf-binding sites, whereas the *dkk1* and *lhx5* enhancers have both Sox17 and Tcf sites. To test whether or not Tcfs regulate these enhancers, we injected a Tcf7l1-VP16 fusion construct which constitutively activates Tcf-target gene transcription (Darken and Wilson, 2001). As predicted, this had little to no impact on the *six1* reporter, but robustly activated the *dkk1* and *lhx5* reporters in ventral ectoderm (Fig. 6D,F, S7B,D,G). We next mutated the Sox17 and Tcf binding sites in the *six1* and *dkk1* enhancers and examined the impact on Sox17, Bcat and Tcf regulation. Loss of the Sox17 sites abolished normal and Sox17/Bcat-induced expression of the *six1:luc* reporter (Fig. 6D). In contrast mutation of the Sox sites in the enhancers resulted in elevated reporter activity consistent with Sox17-mediated repression (Fig. 6F and S7G). Deletion of the Tcf sites in the *dkk1* enhancers abolished expression in the dorsal mesendoderm as well as Bcat or Tcf7l1-VP16 induced activation in the ventral ectoderm, consistent with Bcat-Tcf complexes activating these enhancers (Fig. 6F and S7G).

From these collective data, we conclude that Sox17 and Bcat synergistically activate the *six1* enhancer, likely in a Tcf-independent manner. In contrast, the *dkk1 and lhx5* enhancers are activated through a Bcat/Tcf transcription mechanism, with Sox17 spatially restricting Bcat/Tcf activity to maintain cell-type specific gene expression (Fig. 7).

**Fig 7:**
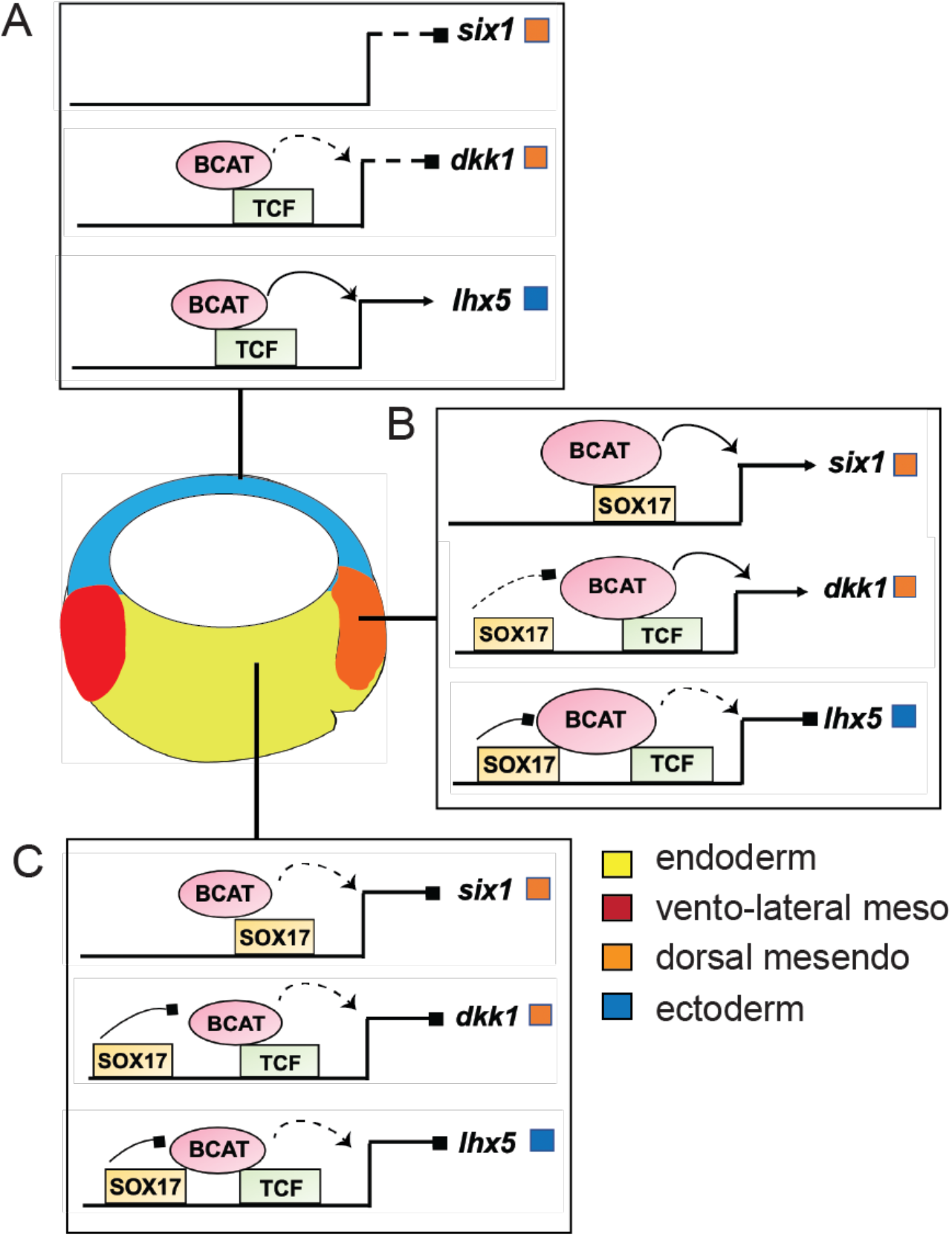
Model of Sox17 and Bcat coregulation spatial trasncription: Differential engagement of Sox17, Bcat and Tcfs on enhancers in different cells modulates spatial expression domains. **(A)** In ectoderm cells lacking Sox17 (and Nodal signaling), ectoderm-specific gene *lhx5* is activated through a Bcat/Tcf dependent mechanism while *six1* and *dkk1* are not transcribed. (**B-C**) Sox17 is expressed throughout the endoderm and dorsal mesendoderm, while Bcat activity is the higher in dorsal mesendoderm than deep endoderm. **(B)** In the dorsal region Sox17 and Bcat synergistically coactivate *six1* in the absence of Tcf, whereas Sox17 exerts a repressive indluence on Bcat/Tcf-activation of *lhx5 and dkk1*. We postulate however that in the dorsal region Sox17-mediated repression is insufficient to overcome Bcat/Tcf activation of *dkk1*. **(C)** In deep endoderm cells where Bcat activity is lower, Sox17 mediated repression of Bcat/Tcf is sufficient to repress *dkk1* and lhx5 transcription, but Bcat activity is not high enough to activate *six1*.

## DISCUSSION

### Overview of findings

Sox17 is a key regulator of vertebrate endoderm development and morphogenesis, yet surprisingly, the transcriptional program that it regulates has been poorly characterized until now. Here we defined the Sox17-regulated GRN in *Xenopus* gastrulae, which includes many conserved SOX17 genomic targets in PSC-derived human definitive endoderm. We show that in addition to acting atop a transcriptional hierarchy promoting endoderm differentiation, Sox17 has previously unappreciated roles in endoderm patterning and germ layer segregation. We demonstrate that functional interaction with the canonical Wnt signaling pathway is a major feature of the Sox17-regulated GRN. Over a third of all Bcat and Sox17 genomic binding events in the gastrula occur at the same enhancers, where they coordinately regulate spatial gene expression in the embryo. Our data suggest that Sox17 and Bcat synergistically activate a subset of endodermal enhancers in the absence of Tcfs, whereas at other enhancers, Sox17 modulates Bcat-Tcf mediated transcriptional activation. Together these results provide novel insights into the establishment of the endoderm GRN and suggest a new paradigm where Sox17 acts as a tissue specific modifier of Wnt-signaling responses.

### The Sox17-Bcat regulated endodermal GRN

Our genomic study revealed an extensive overlap between Sox17- and Bcat-regulated genes. Together with previous work (Charney et al., 2017b; Zorn and Wells, 2009), this suggests that Sox17 and the other core endoderm TFs interact with Wnt and Nodal in a series of positive and negative feedback loops to control germ layer segregation and modulate spatial gene expression. We showed that in addition to its known role in activating transcription of endoderm-inducing *nodal* ligands (Blythe et al., 2010; Hyde and Old, 2000), Bcat directly promoted the expression of many endodermal genes including most of the core endoderm TFs; *sox17, eomes, foxa, gata4/5, mix1* and *mixer*. Sox17 also regulated the expression of other endoderm GRN components; promoting expression of *foxa1* and *gata4*, whilst negatively regulating the expression of some *nodal* and *mix*/*bix* genes. Thus on one hand, Sox17 promotes endodermal fate, while it restrains excessive Nodal signaling on the other. There is precedence for Sox factors influencing *nodal* ligand transcription. Maternal Sox3, localized in the blastula ectoderm, directly binds the *nodal5/6* enhancers to repress Bcat-stimulated transcription, whereas vegetally localized maternal Sox7 enhances *nodal5/6* expression in the vegetal cells (Zhang and Klymkowsky, 2007). This suggests a potential handoff model where different Sox TFs might co-occupy the same enhancers with Bcat in different cells to activate or repress transcription in a context-dependent manner.

Our data demonstrate that Sox17-Bcat interactions regulate spatial gene expression in the embryo (Fig. 7). Sox17-Bcat interact at some enhancers like *six1* and *osr1* to promote transcription in the endoderm and dorsal mesendoderm. In addition, Sox17 also has a key role in repressing alternative lineages in the vegetal cells, similar to the maternal TFs Foxh1, Vegt and Otx1 (Chiu et al., 2014; Paraiso et al., 2019). For example, Sox17 represses Bcat/Tcf-stimulated expression of the ectoderm specifying TFs *lhx5* (Houston and Wylie, 2003). Sox17-Bcat interactions also impact spatial expression within the endoderm. Sox17 inhibits the Bcat-Tcf stimulation of the *dkk1* enhancer in the deep endoderm, but in the dorsal mesendoderm where Bcat activity is higher, Sox17 appears insufficient to suppress *dkk1*. The finding that *six1* and *dkk1* are under opposite regulation by Sox17 and Bcat despite having very similar expression patterns warrants future analysis of the temporal-spatial dynamics in Bcat, Tcf and Sox17 genomic occupancy in dorsal vs ventral cells over time.

### DNA Binding Specificity of SOX TFs

An important consideration is how Sox17 selects specific target enhancers. In DNA-binding experiments, all Sox TFs bind to variants of the DNA sequence 5’-(A/T)(A/T)CAA(A/T)3’, which is similar but distinct from Tcf DNA-binding sites 5’-T(A/T)(A/T)CAAG 3’ (Hou et al., 2017; She and Yang, 2015). SOX TFs can act either as monomers or heterodimers with different cell-type specific cofactors, which in turn impact their DNA-binding affinity and ability to activate or repress transcription (Bernard and Harley, 2010; Hou et al., 2017). For example, in PSCs, Oct4-Sox2 heterodimers promote pluripotency and bind to distinct sites than those bound by Oct4-Sox17 complexes that initiate mesendoderm development (Aksoy et al., 2013). It is, therefore, likely, that Sox17 and Bcat cooperate with other TFs to provide lineage specificity to Wnt-responsive transcription during germ layer segregation.

Motif analysis revealed that Sox17 bound loci were also enriched for Tbx and Gata motifs indicating that in addition to regulating each other’s expression, the core endoderm TFs act together in combinatorial manner, perhaps to establish endodermal super enhancers crucial for downstream lineage specification (Paraiso et al., 2019). Interestingly, enhancers that were negatively regulated by Sox17 were enriched for Tbx and Oct motifs whereas enhancers activated by Sox17 were enriched for homeodomain motifs. This suggests that Sox17 interactions with distinct lineage-specific TFs could elicit a context-specific transcriptional response. In the future, it will be important to identify and integrate the genomic targets of all the core endoderm-specifying TFs and examine how combinatorial enhancer binding is integrated into a unified endodermal GRN.

### Sox-Bcat-Tcf Interactions on chromatin

Many SOX proteins, including Sox17, can bind to recombinant Bcat and Tcfs *in vitro* and modulate Wnt/Bcat-stimulated transcription of synthetic reporters assays with multimerized Tcf-sites (pTOPFlash) (Sinner et al., 2007; Zorn et al., 1999). While a number of models have been proposed to explain how Sox factors might impact Bcat activity, (Kormish et al., 2010; She and Yang, 2015; Tan et al., 2019), in most cases the *in vivo* biological relevance of these interactions was unclear. Although the resolution of ChIP-reChIP assays only allow us to demonstrate that Sox17 and Bcat bind chromatin within 150bp of each other, taken together with published *in vitro* data, our results suggest that Sox-Bcat or Sox-Bcat-Tcf transcriptional complexes can be assembled on enhancers *in vivo*.

At enhancers like *six1* where Sox17 and Bcat synergistically stimulate transcription, we postulate that Sox17 and Bcat recruit Ep300/Cbp to chromatin, analogous to Tcf-Bcat in canonical Wnt-signaling. Consistent with this, structure-function studies have found that a conserved 10 amino-acid domain (VDXXEFEQYL) in the Sox17 C-terminus is required for both Bcat-binding and transcriptional transactivation (Sinner et al., 2004). On the other hand, most Bcat and Sox17 co-occupied enhancers contain multiple Sox and Tcf sites clustered within 50 bp, suggesting that Sox17 and Tcf, together with Bcat and scaffolding proteins like Bcl9, (Gammons and Bienz, 2018; Masuda and Ishitani, 2017) might assemble higher-order multiprotein complexes facilitating Wnt-mediated transcription. Both Soxs and Tcfs are HMG-domain containing TFs, capable of binding to the minor groove of DNA and inducing DNA bending. This could further indicate that they are brought into close spatial proximity to establish the assembly of functional enhanceosomes, despite the distance between their binding sites on linear DNA (Bernard and Harley, 2010; Hou et al., 2017).

The possibility of Sox-Bcat-Tcf ternary complexes raises the question of competitive vs cooperative DNA binding that, in turn, could influence transcriptional repression versus activation. For example, Sox17 might stabilize the formation of Bcat-Tcf-Ep300 complexes or destabilize the binding of co-repressors such as Tle and Hdac to promote transcription (e.g., *osr1*). To negatively regulate Bcat-Tcf activity (e.g.,*dkk1*) Sox17 might destabilize Bcat-Tcf-Ep300-DNA or stabilize Tcf-Tle complexes. This idea is supported by work in mouse cortical neuron progenitors where Sox2 binds within 4-10bp of Tcf/Lef sites and recruits Tle/Gro corepressors to repress Tcf/Bcat-mediated *Ccnd1* expression (Hagey and Muhr, 2014). Future biochemical studies will need to determine if and how Sox-Bcat-Tcf complexes assemble on DNA.

### Broader Implications

Although a growing number of non-Tcf Bcat-binding TFs have been reported, (Gammons and Bienz, 2018; Notani et al., 2010; Trompouki et al., 2011), how these impact the genomic specificity of Wnt-responsive transcription has not been well characterized and the idea of functional Tcf-independent Wnt transcription remains controversial (Doumpas et al., 2019; Schuijers et al., 2014). Our data suggest that in the gastrula endoderm, Sox17 might influence Bcat recruitment to chromatin to control cell-type specific Wnt-responsive transcription, and in some cases, this might could occur independent of Tcfs. Given that there are 20 Sox TFs encoded in the vertebrate genome with most cell types expressing at least one, it is possible that SOX factors are widespread and previously unappreciated accessory effectors of Bcat that finetune Wnt-regulated transcription across many biological contexts.

## METHODS

### *Xenopus* embryo manipulations

*Xenopus laevis* and *Xenopus tropicalis* were purchased from Nasco or the *National Xenopus Resource* and experiments were performed according to CCHMC IACUC approved protocols. Embryos were staged according to Nieuwkoop and Faber (NF) (Nieuwkoop and Faber, 1967). Microinjection and embryo culture were preformed as previously described (Stevens et al., 2017) Antisense morpholino oligos (MO; GeneTools, LLC) were as follows:

Standard control MO: 5’CCTCTTACCTCAGTTACAATTTATA-3’;
*X.tropicalis ß-catenin* MO: 5’ TTTCAACAGTTTCCAAAGAACCAGG 3’;
*X. tropicalis sox17a* MO: 5’-AGCCACCATCAGGGCTGCTCATGGT-3’
*X. tropicalis sox17b.1/2* MO: 5’-AGCCACCATCTGGGCTGCTCATGGT-3’
*X.laevis ß-catenin* MO: 5’TTTCAACCGTTTCCAAAGAACCAGG-3’ (Heasman et al., 2000);
*X.laevis sox17a* MO: 5’-ATGATGAGGAGGGTTGGACAGGAGA-3’ (Clements et al., 2003);
*X.laevis sox17b1/2* MO: 5’-TGATTCGGAGTGCTGTGGTGATTAG-3’ (Clements et al., 2003);
Total amounts of MO injected into *X.tropicalis* embryos were as follows: 2ng Bcat MO, 8ng sox17 MO (4ng each a+b1/2 MO), 8ng standard control MO.

Synthetic mRNA for injections was generated using the Message Machine SP6 transcription kit (Thermo Fisher AM1340) using the following plasmids: pCS2+ human B-catenin S37A caBCAT) (Zorn et al., 1999); pcDNA6-V5-mouse Sox17 (mSox17) and pcDNA6-V5-Sox17 M76A (Sinner et al., 2007); pCS2+ Tcf7l1:VP16 (formerly XTcf3:VP16) (Darken and Wilson, 2001).

### *In situ* hybridization

In situ hybridization of *Xenopus tropicalis* embryos was performed as described (Sive et al., 2000) with the following minor modifications. After overnight fixation at 4°C in MEMFA, embryos were washed (2×10 minutes) in MEMFA buffer without the formaldehyde and stored in 100% ethanol at −20°C. Embryos were serially rehydrated to PBS+0.01% Tween-20 and bisected through the dorsal-ventral axis in on 2% agarose coated dishes using a fine razor blade (Electron Microscopy Services 72003-01), followed by Proteinase K (ThermoFisher AM2548) treatment at 1ug/mL for 10 minutes. TEA/Acetic Anhydride steps were skipped on day1. The RNAse A step was 0.25ug/mL for 5 minutes on day 2; and finally the anti-DIG-alkaline phosphatase antibody (Sigma 11093274910) was used at a 1:6,000 dilution in MAB buffer + 20% heat-inactivated lamb serum + 2% blocking reagent (Sigma 11096176001) on day2/3. Anti-sense DIG labeled in situ probes were generated using linearized cDNA templates with the 10X DIG RNA labeling mix (Sigma 11277073910) according to manufacturer’s instructions. Details on plasmids used for probe synthesis are listed in supplementary table S7.

### Antibodies and Immunostaining

Affinity purified anti-*Xenopus* Sox17 rabbit polyclonal antibodies were generated by Bethyl labs to the Sox17a/b N-terminal, Sox17b C-terminal and Sox17a C-terminal with the peptides illustrated in Fig. S2. Antibodies were validated by immunostaining, Western blot, immunoprecipitation and ChIP. Immunoreactivity was not detected in Sox17 deficient tissue and specifically competed by addition of the target peptide to reactions. In preliminary ChIP-qPCR experiments, the pan-Sox17 and Sox17bC-terminal antibodies bound to the same loci, but the Sox17bC-terminal was more efficient and was used for ChIP-seq.

For *Xenopus* immunofluorescence, embryos were fixed in ice-cold 100mM NaCl, 100mM HEPES (pH7.5), 3.7% methanol-free paraformaldehyde for 1.5 hours at 4°C and then dehydrated directly into ice-cold Dent’s post-fixative (80%Methanol/20% DMSO) and stored at −20°C. Embryos were serially rehydrated on ice into PBS+0.1% TritonX-100 (PBSTr) and bisected through the dorsal-ventral axis with a fine razor blade. Embryo halves were subjected to antigen retrieval in 1x R-Universal epitope recovery buffer (Electron Microscopy Services #62719-10) for 30 min at 55-60°C, washed 2 × 10 minutes in PBSTr, blocked for 2 hours in PBSTr + 10% normal donkey serum (Jackson Immunoresearch 017-000-001) + 0.2% DMSO at room temperature, and then incubated overnight at 4°C in this blocking solution + the following primary antibodies: chicken anti-GFP (Aves GPF-1020; diluted 1:1,000), rabbit anti-Sox17a/b N-terminal (1:350), rabbit anti-B-catenin (Santa Cruz Biotechnology sc-7199; 1:500). After extensive washing in PBSTr, samples were incubated overnight at 4°C in PBSTr + secondary antibodies: donkey anti-Chicken 488, donkey anti-rabbit Cy3 (Jackson ImmunoResearch # 703-546-155, #711-166-152, both used at 1:1,000 dilution).TO-PRO3 nuclear stain (ThermoFisher #T3605) was included in one 30min PBSTr wash after secondary antibody incubation. Samples were extensively washed in PBSTr, dehydrated into 100% methanol, cleared and imaged in Murray’s Clear (2 parts benzyl benzoate, 1 part benzyl alcohol).

### Luciferase Assays

Putative enhancers (wildtype as Sox and Tcf-site mutants) were synthesized (supplementary sequence file S1) and cloned into the pGL4.23 *luc2*/miniP vector. (Promega E8411). *Xenopus laevis* embryos were used for luciferase assays because of the larger size embryos and slower developmental rate. Enhancer activity was assayed by co-injecting embryos with 5pg of pRL-TK-renilla luciferase vector (Promega E2241), 50pg of the pGL4.23 *luc2*/miniPenhancer:luciferase and combinations of the following MOS or mRNAs: 5pg ca-Bcat, 25pg mSox17, 25pg mSox17 M76A, 10pg Tcf3-VP16, 20ng *Xl sox17a+b1/2* MO (10ngeach), 10ng *Xl bcat* MO. For each condition three tubes of 5 NF10.5 embryos were collected. Embryos were lysed in 100uL of 100mM TRIS-Cl pH7.5, centrifuged for 10 minutes at ~13,000 x *g* and then 25uL of the clear supernatant lysate was used separately in Firefly (Biotium #30085-1) and Renilla (Biotium 30082-1) luciferase assays according to the manufacturer’s instructions. The average relative Luciferase activity was normalized Renilla levels for each sample and statistical significance was determined by pairwise Student T-test.

### RNA-Sequencing

#### RNA isolation and library preparation

Approximately 200 *X. tropicalis* embryos were injected with 8 ng of either control-MO or Sox17-MO, or 2 ng of control-MO and Bcat-MO and 10 sibling embryos were collected at every time point from blastula to late gastrula in biological duplicate. Total RNA was extracted with the Nucleo-spin RNA kit (Machery-Nagel). RNA-seq libraries were generated using Smart-seq2 cDNA synthesis followed by tagmentation, quality-tested using an Agilent Bioanalyzer 2100, quantified using KAPA qPCR and sequenced using Illumina sequencers at the UC Irvine Genomics High Throughput Facility.

#### RNA-Seq Analysis

Raw reads were quality checked using FastQC and adapters/low quality reads were removed with Trimmomatic (Bolger et al., 2014). Reads were mapped to the *Xenopus tropicalis* genome version 9.0 (Hellsten et al., 2010) with bowtie2, quantified using RSEM (Li and Dewey, 2011) and reported in transcripts per million (TPM). *X. tropicalis* v9.0 gene name annotations were obtained from Xenbase.org (Karimi et al., 2018). Differentially expressed genes (≥2 fold change and FDR ≤ 5%) were identified by a pairwise comparison between control-MO versus Sox17-MO and control-MO versus Bcat-MO sibling embryos at each stage using RUVSeq (R package) (Risso et al., 2014). Gene ontology enrichment analysis was performed using the Gene Ontology Consortium Online Resource (http://geneontology.org).

To assess the spatial expression of transcripts we examined the TPM expression values in dissected stage 10.5 embryos from published data in GEO GSE81458 (Blitz et al., 2017). Enriched expression in a specific tissue was defined as the normalized expression being >1.5x of the mean expression across dissected tissues. Statistical significance of enriched expression pattern in groups of genes was determined by chi-square test p<0.05 performed in R package MASS.

### Chromatin Immunoprecipitation

#### ChlP-Seq

Large scale Sox17 ChIP was carried out as previously described (Charney et al., 2017a) using 2000 NF10.5 *X. tropicalis* embryos and 30 μg rabbit anti-Sox17b-Cterminal antibody. ChIP-seq libraries were generated using Nextflex ChIP-seq kit (Bio Scientific), analyzed using an Agilent Bioanalyzer 2100, quantified using KAPA qPCR and sequenced using Illumina instruments at the UC Irvine Genomics High Throughput Facility.

#### ChlP-qPCR

was carried out largely as described in (Charney et al., 2017a; Hontelez et al., 2015) with minor modifications. Briefly, 50-200 NF 10.5 *Xenopus tropicalis* embryos were fixed in 1% formaldehyde in 0.1x MBS at room temperature for 30 minutes. Sonication was carried out in a Diagenode Bioruptor Pico for 30 cycles of 30 seconds ON, 30 seconds OFF. Chromatin extracts were incubated with 10 μg rabbit anti-Sox17-Cterminal antibody (affinity purified by Bethyl labs, FigS2) or 2.5 μg rabbit anti-β-catenin (Life Technologies, 712700) overnight at 4°C. After washes and elution, DNA was then purified using a MinElute PCR Purification kit and QIAQuick purification columns and eluted in 20μl TE.

#### ChlP-reChIP

Before the first ChIP, the anti-β-catenin antibody was crosslinked to Protein G Dynabeads. ChIP assay was then carried out on extracts from 500 embryos as described above. At the end of the first ChIP, DNA was eluted with elution buffer supplemented with 10mM DTT. The eluate was then diluted in 10 volumes of wash buffer (50mM Tris-HCl pH 8.0, 100mM NaCl, 2mM EDTA, 1% NP-40) supplemented with 1x Protease Inhibitor Cocktail and 1mM DTT. The 2^nd^ ChIP was then carried out as a normal ChIP as described above.

#### ChlP-Seq Analysis

Raw reads were assessed by FastQC and trimmed with Trimmomatic and aligned to *Xenopus tropicalis* genome (Hellsten et al., 2010) version 9.0 using bowtie2 v2.3.4.7 (Langmead and Salzberg, 2012). Duplicate and multi-mapped reads were removed by Picard (http://broadinstitute.github.io/picard/) and samtools (Li et al., 2009). Peaks were called with MACS2 (Zhang et al., 2008) against NF10.5 input DNA with default options. Peaks were associated with genes using the HOMER nearest TSS function. To identify statistically significant ChIP-seq peaks we performed Irreproducible discovery rate (IDR) analysis per ENCODE Best Practices guidelines (Landt et al., 2012) on pseudoreplicate from pooled Sox17 ChIP-seq biological replicates and pseudoreplicate of the single Bcat ChIP-seq dataset from GEO GSE72657 (Nakamura et al., 2016) with an IDR threshold of 0.01. The location of peaks (intron, intergenic, exon and promoter) were annotated using HOMER v4.1.0 (Heinz et al., 2010). To visualize the data, aligned bam files generated as above were converted to bigwig format using deeptools v3.4.1 and visualized on IGV v2.4.1.3 (Thorvaldsdottir et al., 2013). Significance of intersected ChIP-seq and RNA-seq datasets was assessed using hyper geometric test (HGT).

#### Motif analysis

*De-novo* motif analysis was performed with HOMER v4.1.0 (Heinz et al., 2010). For a comprehensive Sox and Tcf motif analysis, we divided our datasets into 3 groups: a) Peaks bound and regulated by Sox17 only, b) Peaks bound and regulated by Bcat Only and c) Peaks that were cobound and coregulated by Sox17 and Bcat. DNA sequences of 150 bp centered on the peaks summits were extracted and motif analysis was carried out as follows: 1) To find which peaks have instances of Sox17 and Tcf motifs, HOMER function annotatePeaks.pl was used with the parameters: −size 150 −m motif sox17.motif tcf7.motif lef1.motif tcf3.motif tcf7l2.motif. 2) To perform a more exhaustive motif search the enhancers were also scanned for presence of position weight matrices (PWMs) of experimentally defined Sox17, Tcf7, Tcf7l1 and Lef1 motifs > 70% threshold from the CIS-BP database (Lambert et al., 2019). The HOMER and CIS-BP results were merged to generate BED files and Bedtools v2.27.1 (Quinlan and Hall, 2010) intersect function was then used to extract coordinates of ‘Sox only’ ‘Tcf only’ and ‘Sox and Tcf overlapping’ categories.

#### Peak density heatmaps

From the Sox17 and Bcat peaks called by MACS2, Bedtools intersect function was used to stratify the peaks into ‘Sox bound only’, ‘Bcat bound only’ and ‘Sox and Bcat overlapping peaks’. For each category of peaks, the peak summit was extracted from the TF summit file generated by MACS2 and 2000 bp were added to both sides of the summit using awk. Bedtools computematrix was used to generate per genome regions. The generated matrix was then used with bedtools plotHeatmap to generate heatmaps and signal density plots.

### Analysis of public datasets

The following datasets were downloaded from GEO: Embryo dissection RNA-seq; GSE81458 (Blitz et al., 2017), embryonic stage series RNA-seq; GSE65785; (Owens et al., 2016), Bcat ChIP-seq; GSE72657 (Nakamura et al., 2016), RNA pol II ChIP-seq; GSE85273 (Charney et al., 2017a), H3K27Ac ChIP-seq; GSE56000 (Gupta et al., 2014), H3K4me1 and Ep300 ChIP-seq; GSE67974 (Hontelez et al., 2015). Wherever necessary, raw data was downloaded from GEO and aligned to *Xenopus* genome version 9.0, and processing to generate bigwig files and heatmaps was carried out as described above.

### Data and software availability

The RNA-seq and ChIP-seq data generated by this study have been deposited in the NCBI Gene Expression Omnibus (GEO) and can be accessed under GSE148726 (reviewer password: choluuuefpkprsb).

## Supporting information

Supplemental Table1_Sox17_Metadata

Supplemental Table2_Sox17_ChIP_peaks

Supplemental Table3_Bcat_Metadata

Supplemental_Table4_Bcat_ChIP_peaks

Supplemental Table5_Cobound_Coreg_enhancers

Supplemental Table6_ChIP_Primers

Supplemental enhancer sequences

## Acknowledgements

This work was supported by HD073179 to AMZ and KWYC and by NIH P30 DK078392 gene expression core of the Digestive Diseases Research Core Center in Cincinnati. We thank the University of California (UC) Irvine Genomic High-Throughput Facility Shared Resource of the Cancer Center Support Grant (P30CA-062203) and the UC Riverside Institute for Integrative Genome Biology for sequencing support. We are grateful to members of the Zorn and Wells labs and the Endoderm Club for helpful discussions. We also wish to thank Xenbase.org (RRID: SCR_003280) and the National *Xenopus* Resource (RRID:SCR_013731) for critical community resources.

## Author Contributions

AMZ, KWYC, ILB, MBF, SAR and SM designed the experiments. SAR, MBF, MW, MM, SM and ILB performed the experiments. SM, PC, KDP, XC and MTW performed the bioinformatic analysis. SM, SAR, ILB, KWYC and AMZ wrote the manuscript.

---

**Fig. S1.**
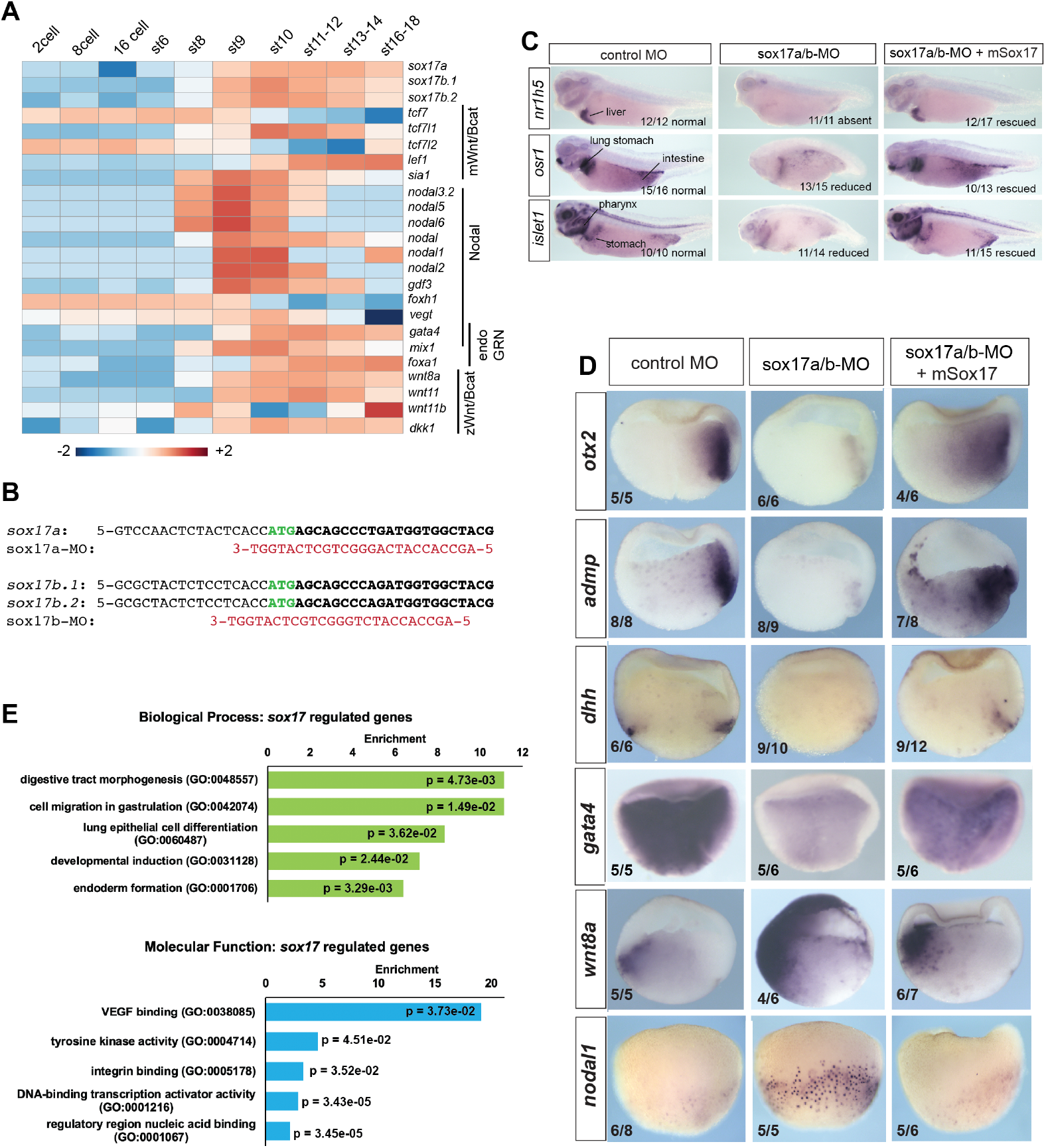
Characterization of Sox17-MO embryos and Sox17-regulated transctiptome. **(A)** Temporal expression of key endoderm GRN transcripts (GSE65785; (Owens et al., 2016). **(B)** Sequence alignment of antisense sox17a-MO and sox17b-MO (red) with 5’ end of *Xenopus tropicalis sox17a, sox17b.1* and *sox17b.2* mRNAs (black) spanning the ATG translation start (green). **(C)** In situ hybridization of markers for liver (*nr1h5*), stomach, lung, intestine and pharynx (*osr1* and *islet1*) snowing that co-injection of mouse *Sox17* RNA rescues gut development in sox17-MO embryos at the tailbud stage. **(D)** Additional validation of gastrula stage trasncripts downregulated (*otx2, admp, dhh* and *gata4*) or upregulated (*wnt8* and *nodal1*) in sox17-MO embryos and resuce by co-injection of mSox17. **(E)** GO term enrichment analysis of Sox17-regulated genes.

**Fig. S2.**
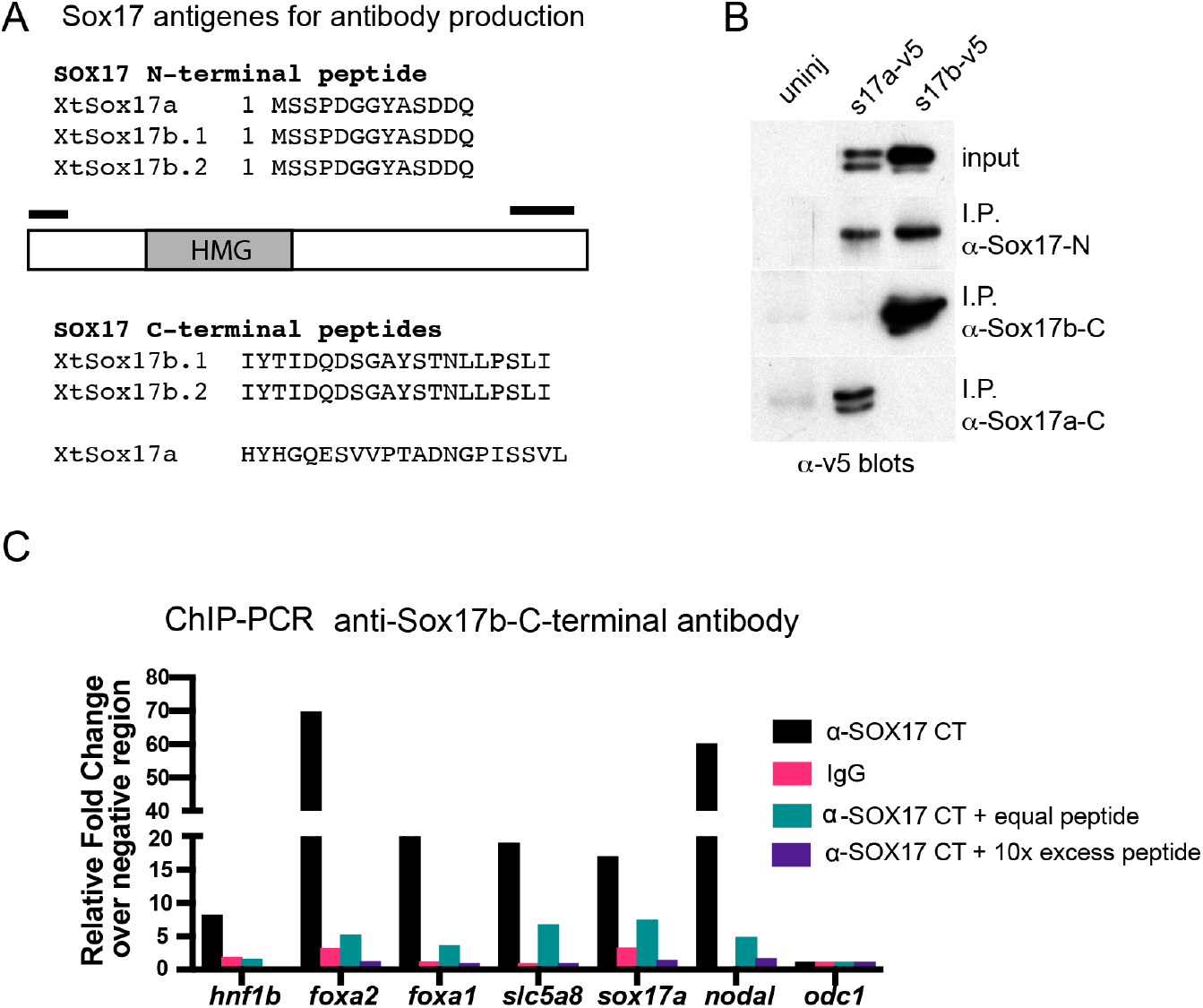
Validation of anti-Sox17 antibodies. **(A)** Schematic and amino acid sequences of the *Xenopus tropicalis* Sox17 peptides uses to immunize rabbits and generate polyclonal antibodies to; pan-Sox17 N-terminal, Sox17b C-terminal and Sox17a C-terminal. **(B)** Immunoprecipitation of *Xenopus laevis* embryos expressing V5 epitope tagged versions of Sox17a-V5 and Sox17b1-V5. Anti-V5 western blot showed that all antibodies specifically immunoprecipitated the target antigen but that the Sox17b C-terminal antibody was the most efficient. **(C)** ChIP-PCR validation of anti-Sox17 antibodies. In preliminary ChIP-PCR experiments the pan-Sox17 N-terminal and Sox17b C-terminal antibodies bound to all the same positive control loci, but the Sox17b C-terminal antibody was more efficient (data not shown) and was used for further optimization and the ChIP-seq analysis. The histogram shows the relative fold change of Sox17b binding versus IgG control in ChIP-PCR of positive control loci and the negative control loci *odc* all normalized to a second negative control loci *eef1a1*. Addition of the Sox17b C-terminal peptide reduced the ChIP signal in a dose-dependent manner indicating specific binding.

**Fig. S3.**
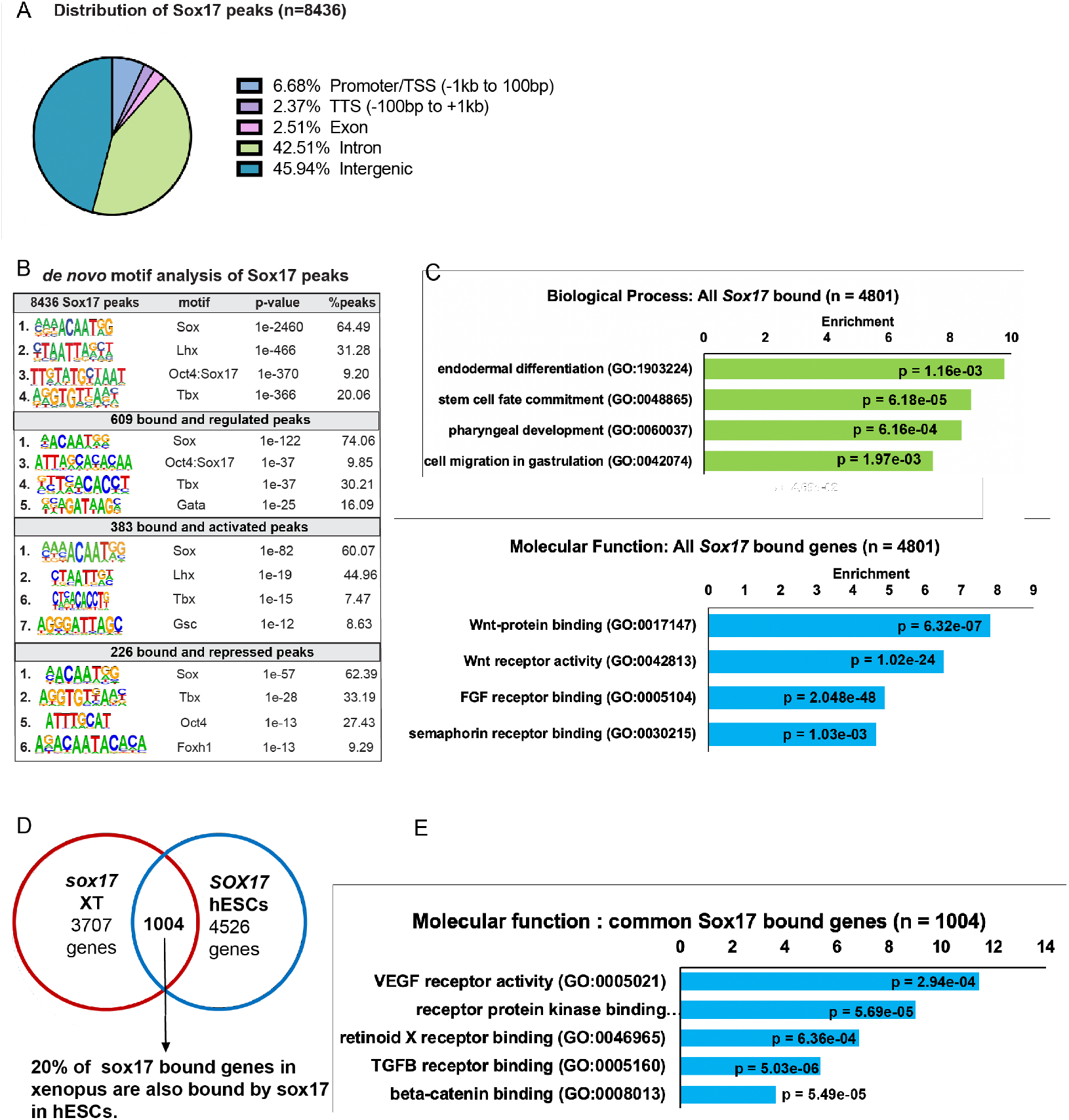
Analysis of Sox17-bound chromatin loci. **(A)** Genomic distriputions of 8436 Sox17 bound chromatin loci. ChIP-seq peaks were associated to cognate promoters using Homer nearest gene transcription start site (TSS). 88% of the Sox17 peaks were located in either introns or elements more than 1 kb from the TSS, indicative of distal CRMs. **(B)** *De novo* motif analysis (Homer) of Sox17 peaks associated with Sox17-regulated genes. **(C)** GO analysis of 4810 genes associated with Sox17 ChIP-seq peaks showed enrichment for endoderm development and Wnt signaling. **(D)** Comparison of the *Xenopus* genes associated with Sox17 ChIP-seq peaks to published human SOX17 ChIP-seq from human embryonic stem cell (hESC)-derived definitive endoderm (GSE61475; (Tsankov et al., 2015). The extensive overlap indicates that Sox17 regulates an evolutionarily conserved GRN. The venn only includes *Xenopus* genes that have human orthologs, unannotated *Xenopus* genes were excluded from the analysis. **(E)** GO term enrichment analysis of the 1004 Sox17-bound genes conserved between *Xenopus* and human.

**Fig. S4.**
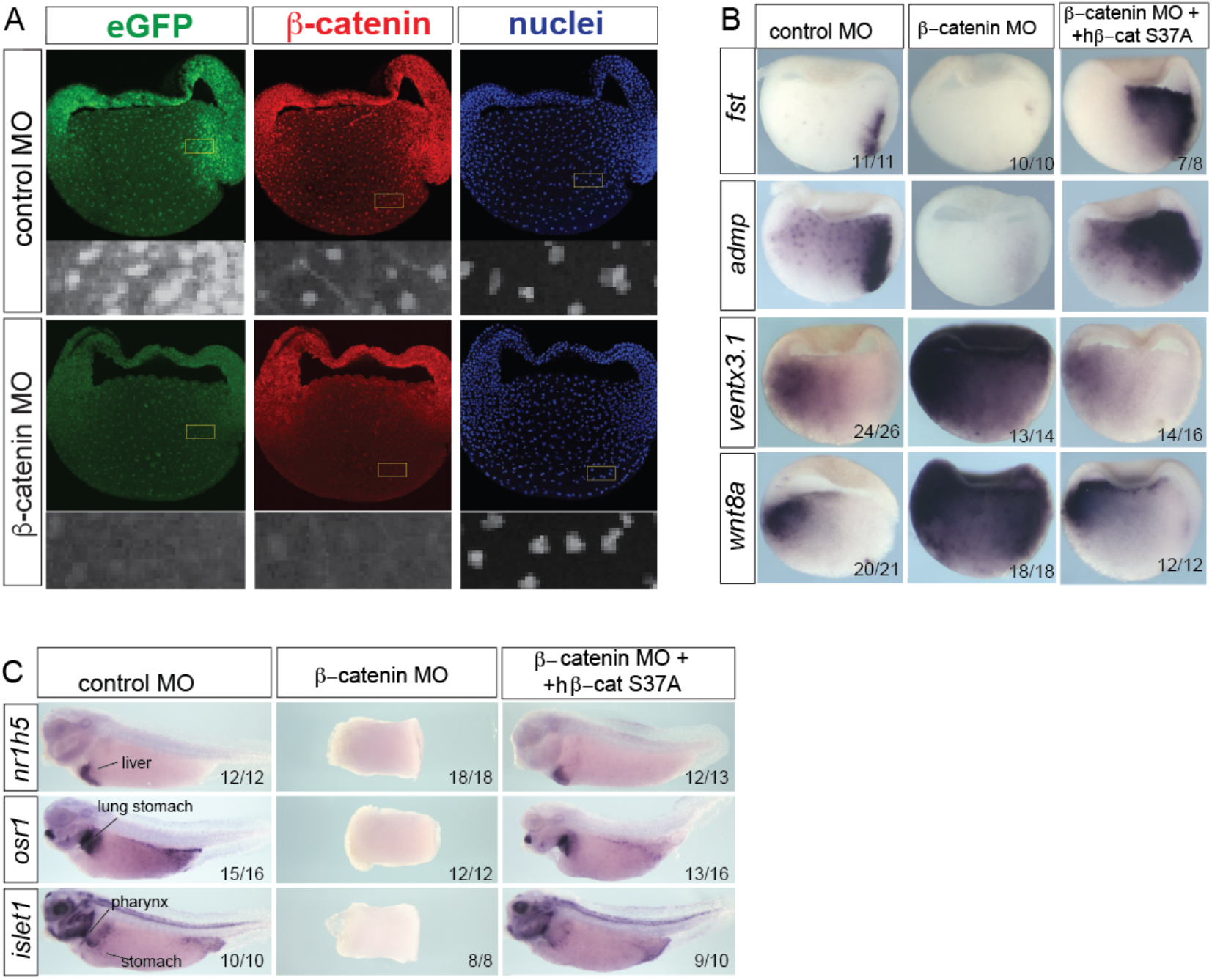
Validation of Bcat-MO embryos. **(A)** Immunostaining of *Tg(WntREs:dEGFP)^Vlem^ Xenopus tropicalis* gastrula shows the loss of nuclear Bcat and GFP expression from the Wnt-reproter transgene in Bcat-MO embryos demonstrating effective knockdown. **(B-C)** Co-injection of RNA encoding a human S37A stabilized Bcat recues the Bcat-MO ventralized phenotype. In situ hybridization of gastrula (B) and tailbud embryos (C). In Bcat-MO gastrula *fst* and *admp* are downregulated in the dorsal organizer,whereas *wnt8a* and *ventx3.1* are upregulated consistent with a ventralized phenotype. (C) In situ hybridization of markers for liver (*nr1h5*), stomach, lung, intestine and pharynx (*osr1* and *islet1*) snowing that co-injection of human S37A stabilized Bcat RNA rescues the ventralized phenotype and disrupted gut development in bcat-MO tailbud embryos.

**Fig. S5.**
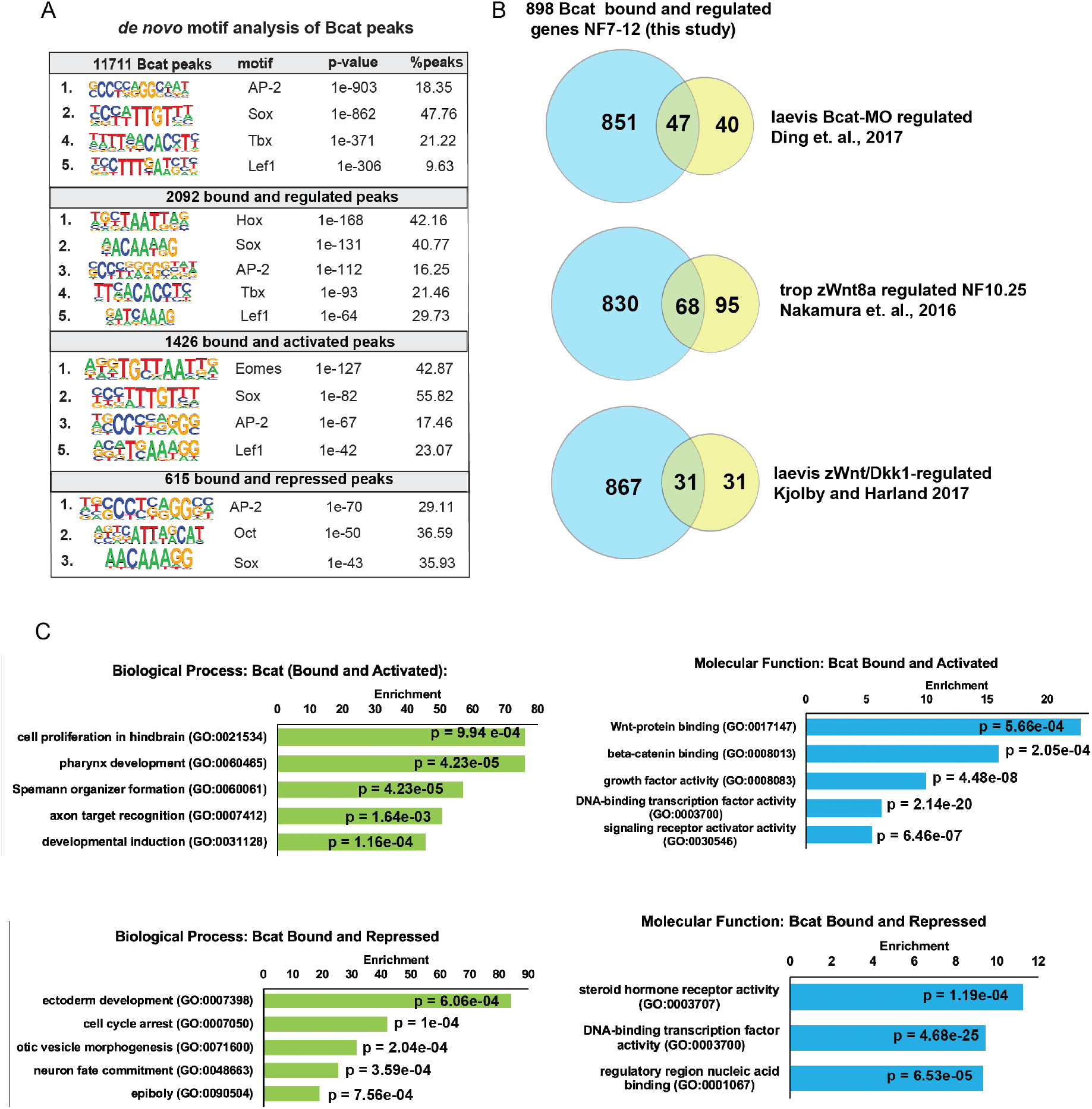
Additional analysis of Bcat ChIP-Seq. **(A)** *de-novo* motif analysis of all Bcat bound, Bcat bound and regulated, Bcat bound and activated or repressed peaks. **(B)** Comparison of differentially expressed genes with other recent genomic analysis of gastrula-stage Wnt targets (Ding et al, 2017; Nakamura et al, 2016; Kjolby and Harland, 2017) **(C)** GO enrichment analysis of peaks corresponding to Bcat Bound and either activated or repressed genes.

**Fig. S6.**
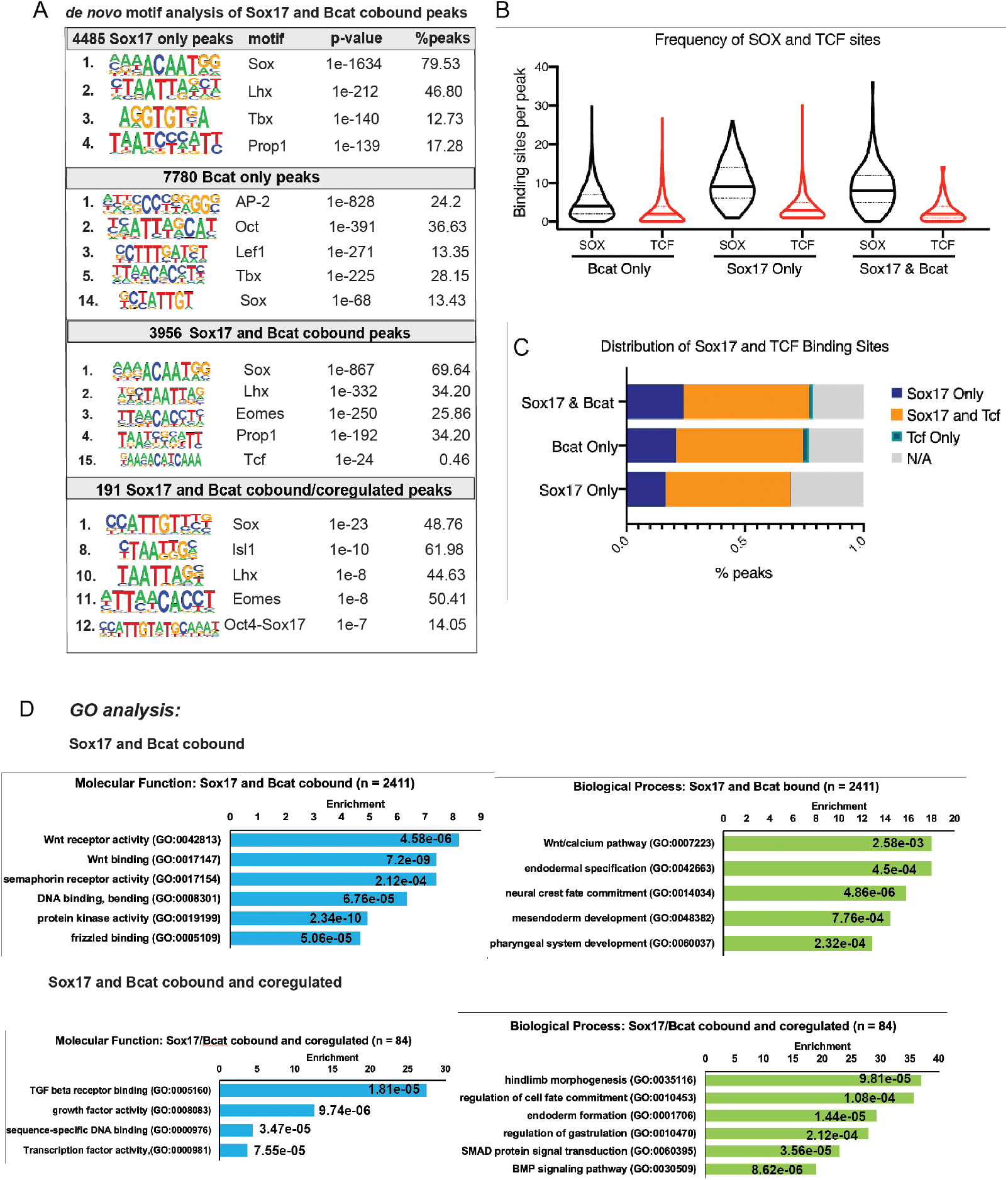
Additional analysis of cobound and coregulated peaks. *de-novo* motif analysis of exclusively bound, cobound and cobound/coregulated Sox17 and Bcat peaks **(B)** Frequency of Sox or Tcf sites per peak in three data subsets: peaks exclusively bound and regulated by Bcat, Sox17 or those that are cobound and coregulated **(C)** Distribution of Sox and Tcf binding sites in three groups of enhancers: those bound and regulated exclusively by Sox17 or Bcat, and those cobound/coregulated by both. **(D)** GO enrichment analysis of cobound and cobound/coregulated Sox17 and Bcat peaks.

**Fig. S7.**
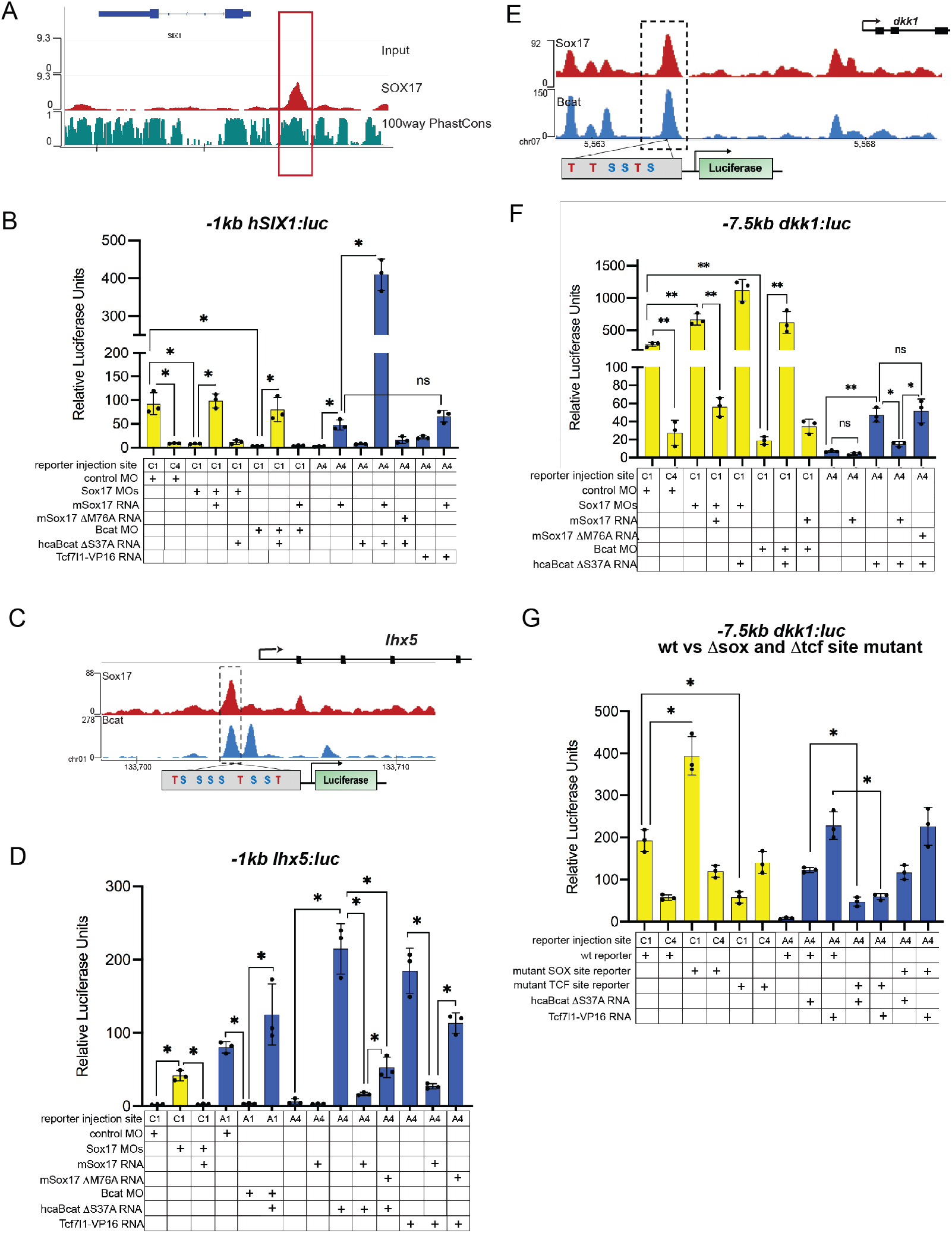
Reporter assays showing Sox17 and Bcat regualted *SIX1*, −7.5kb *dkk1* and −1kb *lhx5* enhancers. **(A)** Browser shot showing evolutionarily conserved *SIX1* peak in human DE (Tsankov et al, 2015). **(B)** Luciferase assay of −1kb *SIX1* enhancer. **(C)** IGV track showing predicted Sox17 and Tcf binding sites at the co-bound *-1kb lhx5* enhancer and **(D)** luciferase assay using the reporter constructed shown in C. **(E)** IGV track showing predicted Sox17 and Tcf binding sites in the −*7.5kb dkk1* enhancer and **(F-G)** luciferase reporter assays of **(F)** the wt −7.5kb *dkk1:luc* and **(G)** wt versus ΔSox-site and ΔTcf-site mutant −7.5kb *dkk1:luc* reporters injected in different tissue with the indicated MOs and/or mRNAs. Standard Deivation, *p<0.05 in pairwise Student T-tests.

## List of Supplementary Data

1. **Supplementary Table S1:** Differentially expressed genes in RNA-Sequencing of Control and and Sox17MO embryos across NF 9-12.
2. **Supplementary Table S2:** Coordinates of Sox17 bound peaks at NF10.5 with their gene annotations.
3. **Supplementary Table S3:** Differentially expressed genes in RNA-Sequencing of Control and BcatMO embryos across NF 9 −12.
4. **Supplementary Table S4:** Coordinates of Bcat bound peaks at NF10.5 with their gene annotations.
5. **Supplementary Table S5:** Coordinates of 191 Sox17 and Bcat cobound/coregulated enhancers and the frequency of Sox and Tcf sites per peak.
6. **Supplementary Table S6: a.** Plasmid and probe synthesis information **b.** ChIP-qPCR primer table
7. **Supplementary File S1:** List of wild-type and Sox/Tcf mutated sequences used for luciferase assays

## Notes

### Competing Interest Statement

The authors have declared no competing interest.

