## Supplemental enhancer sequences for "Sox17 and β-catenin co-occupy Wnt-responsive enhancers to govern the endodermal gene regulatory network"

**Yellow – Sox17**

**Red– Tcf**

1. **-*1kb WT* *six1* enhancer: Chr08:82158915-82159487**

>wt_six1_TSS

AGTGGTTGGTACAACTGAGATCCCTTCCACTACACCTACTTATAACAGGGGAAAGGGGAATTTGATAGGGAGCAATATTGTTCTCTGTCCCATTGGCTGTGGGCACACGTCTGGCCAACTGTGTCTAATTAATGCGAATATCCTTTATTAGGTCATGTGATCATTCTTACCTGATAGCCAAAGTGTATGAAATGACTAGTCAGTCAGTGCCTGAGGATTGCTTTCCCACCTCTCTGTTCCCTTATCTGAACCTATTGTCTTATCTCTGAGCCTCCAGTGCTACCAACAATACATTATCTGTGTTTACAAAGCTAATTGTTTGGGAAATCTTTTTAATAATTGCTTTCAAACAAGGTGTGCAAATGAGGATCAACTTAATTAGAGTGGGACTTCTCTGGGCAATTTTACTAAATACCCTCTTCAAATAAACGAAATATCATGCCTCAGAGTGTTTTCTTTCAGCCTTGCCATTCTCTCTTCAATTAGCCAAGTATTTCCCCACACCCACGGGCCATTATTTCCTCTGGATTCCTGCAAAAGAGGGCAAATTATACGGCTTAACAAAACAAA

2. **-*1kb* ΔSox17 *six1* dSox17: Chr08:82158915-82159487**

>dSox17_six1

AGTGGTTGGTACAACTGAGATCCCTTCCACTACACCTACTTATAACAGGGGAAAGGGGAATTTGATAGGGAGCAATCGCGATCTCTGTCCCATTGGCTGTGGGCACACGTCTGGCCAACTGTGTCTAATTAATGCGAATATCCTTTATTAGGTCATGTGATCATTCTTACCTGATAGCCAAAGTGTATGAAATGACTAGTCAGTCAGTGCCTGAGGATTGCTTTCCCACCTCTCTGTTCCCTTATCTGACCGCGCATCTTATCTCTGAGCCTCCAGTGCTACCAACAATACATTATCTGTGTTTACAAAGCCCGCGCATTTGGGAAATCTTTTTAATAATTGCTTTCAAACAAGGTGTGCAAATGAGGATCAACTTAATTAGAGTGGGACTTCTCTGGGCAATTTTACTAAATACCCTCTTCAAATAAACGAAATATCATGCCTCAGAGTGTTTTCTTTCAGCCTTGCCATTCTCTCTTCAATTAGCCAAGTATTTCCCCACACCCACGGGCCATTATTTCCTCTGGATTCCTGCAAAAGAGGGCAAATTATACGGCTTAACAAAACAAA

3. **Human *-1kb SIX1* enhancer: chr14:61117408-61117704**

>-1kb_six1_hg19

TCATCTGCTGTTTTGTTTATGTGTGGAGAATGCTTAGTAAAACTGCACAAACAAGTGAAGTTGTAATTATCTTAAGCCTCATTTGCACACCTAGTTCCAAAGCAATAATTAAAAGGATTTTGAAAACAATTAGCCTCTCTAAACAAAGATAATCTATTGTCAGTAGCTGAGACTAGCAGGAGATAAGAACAATGCGTGGAGATAAGGGAAGCCAAATGGTGGAAGGAGAGTGACTTTATGACTATACTTATCTCTGATTAGATAAGTTCAAAACAGAATTCTAAGTACACTTCAAAA

4. ***-1.6kb* WT *dkk1_*enhancer: Chr07:5571875-557227**

>wt_dkk1_-2kb

TGAGACCCCCTGGCACAACAACTGATCTATCTATGAAGTCTCCCCCTGACCAATGCCCCCATACATGCACAGAGTGATCAATGCACCTGAGGAAGCAAGATGGCGGCCACAGTAATCACTTGGGCACTGAGTATTTATCACTGCAGCCAATCAAGATGCGAGGAGCTAATGACCAGCGGCCCATTATCACCATCTGATTGTGCCTGTGTCTGGGCTGAGGCCGTTAGAAGGGGGGGACACAGAAGCCCGACACCTTTTGTTCAGTGCAGCCACTAATTGATCTTGGCTAATATAATCAATATTTGATGATCCCGAATCCTTTTCTGCCCCAGACACCTTTTTTATTTGTTATGTGTCcgtacatttacttcatctcttgttttattcttcatatatatag

5. ***1.6kb* ΔSox17 *dkk1_*enhancer (-2kb): Chr07:5571875-557227**

>dSox17_dkk1_-2kb

TGAGACCCCCTGGCTGGATCCTCGATCTATCTATGAAGTCTCCCCCTGACCAATGCCCCCATACATGCACAGAGTGATCAATGCACCTGAGGAAGCAAGATGGCGGCCACAGTAATCACTTGGGCACTGAGTATTTATCACTGCAGCCAATCAAGATGCGAGGAGCTAATGACCAGCGGCCCATTATCCGTATCTAATCTCAGGAGTGTCTGGGCTGAGGCCGTTAGAAGGGGGGGACACAGAAGCCCGACACCTTTTGTTCAGTGCAGCCACTCTTCGGTCTTGGCTAATATAATCAATATTTGATGATCCCGAATCCTTTTCTGCCCCAGACACCTTTTTTATTTGTTATGTGTCcgtacatttacttcagatgacgtcactttcttcatatatatag

6. ***1.6kb* ΔTcf *dkk1_*enhancer: Chr07:5571875-557227**

>dTcf_dkk1_-2kb

TGAGACCCCCTGGCACAACAACTGATCTATATAGTCCAGCTCCCCCTGACCAATGCCCCCATACATGCACAGAGGAGATCTACCACCTGAGGAAGCAAGATGGCGGCCACAGTAATCACTTGGGCACTGAGTATTTATCACTGCAGCCAATCAAGATGCGAGGAGCTAATGACCAGCGGCCCATTATCACCATCTGATTGTGCCTGTGTCTGGGCTGAGGCCGTTAGAAGGGGGGGACACAGAAGCCCGACACCGCCTACGATATGCAGCCACTACTACTCTGTGGCTAATATAATCAATGCCATCACTTCCCGAATCCTTTTCTGCCCCAGACACCTTTTTTATTTGTTATGTGTCcgtacatttacttcatctcttgttttattcttcatatatatag

7. ***-7.5*kb WT *dkk1* enhancer: chr7:5,565,983-5,566,438**

>wt_dkk1 -7.5kb

AGCCGAGCTGCTTGTGATCTCCTTCCCTCCGCTATCAGCATCTCTATTCATTCTGGCTTCAAAGCCGCGGCTTCCCCACACTCATGCCCCTTTCTTCCTCGCCGGTTAGTAATGAAGTTAGGAGCGTGTGTGTTTGCTCGTGCAAACGACATTGTAAGGAAATAGCGGCTATTACAGGCTGAGTGGATTTAGGGAGGCAGAAGCCTTAGAAGTAGGGGGGGGGATTACTCGACCGCTTAACTAGATGAATGTGCTGAGCACAATGTCTGCGCGCCCAAAAGACAACTGAGAAAAGCCGAGCAAACGGACTATTTGTAAAAAGTGGCATCTTTGCTAATGTCTGATTAAAAGCCGGGGTCCCGGGGCTGTTTGATCTGCGCTCATCAAAGGGAGCCTTAAACAAGCTTGGTTTAAAGCCGCTGAAAGGCCTCAGCGCGGGCTGAGAAAACAAAGGCC

8. ***-7.5*kb ΔSox17 *dkk1* enhancer: chr7:5,565,983-5,566,438**

>sox17_dkk1 -7.5kb

AGCCGAGCTGCTTGTGATCTCCTTCCCTCCGCTATCAGCATCTCTATTCATTCTGGCTTCAAAGCCGCGGCTTCCCCACACTCATGCCCCTTTCTTCCTCGCCGGTTAGTAATGAAGTTAGGAGCGTGTGTGTTTGCTCGTGCAAACGACATTGTAAGGAAATAGCGGCTATTACAGGCTGAGTGGATTTAGGGAGGCAGAAGCCTTAGAAGTAGGGGGGGGGATTACTCGACCGCTTAACTAGATGAATGTGCTGAGCGTCCGATCTGCGCGCCCAAAAGACAACTGAGAAAAGCCGAGCAAACGGACGTCCACTAAAAAGTGGCATCTTTGCTAATGTCTGATTAAAAGCCGGGGTCCCGGGGCTGTTTGATCTGCGCTCATCAAAGGGAGCCTTAAAGTCGATTGGTTTAAAGCCGCTGAAAGGCCTCAGCGCGGGCTGAGAAAACAAAGGCC

9. ***-7.5*kb** Δ**Tcf *dkk1* enhancer: chr7:5,565,983-5,566,438**

>sox17_dkk1_-7.5kb

AGCCGAGCTGCTTGTGATCTCCTTCCCTCCGCTATCAGCATCTCTATTCATTCTGGCTTTGCCGCCGCGGCTTCCCCACACTCATGCCCCTTTCTTCCTCGCCGGTTAGTAATGAAGTTAGGAGCGTGTGTGTTTGCTCGTGCAAACGACATTGTAAGGAAATAGCGGCTATTACAGGCTGAGTGGATTTAGGGAGGCAGAAGCCTTAGAAGTAGGGGGGGGGATTACTCGACCGCTTAACTAGAGACCGATGCTGAGCGTCCGATCTGCGCGCCCAAAAGACAACTGAGAAAAGCCGAGCAAACGGACGTCCACTAAAAAGTGGCATCTTTGCTAATGTCTGATTAAAAGCCGGGGTCCCGGGGCTGTTTGATCTGCGCTCATTGCCGGGAGCCTTAAAGTCGATTGGTTTAAAGCCGCTGAAAGGCCTCAGCGCGGGCTGAGAAAACAAAGGCC

10. **TSS WT *lhx5_*enhancer: Chr01:133702140-133702540**

>wt_lhx5_tss

GGGTTACTGGCAGGATTGGTTTGGGCTTTGCTTTACAAAGAAACAGGAGGGGGGTCTGGAATTTTGTTTTCACCTAATCCAGATTTTGTATGTGAATAGGACAGTAGGAGGGAATTCCCTGTAGGACTCCCTAACCACCCCAAACACCAAGGTAAATAAACTTAGCATTAAGCAAACAATTAATACACTGCCCTTTCATAGAACTTCATTATGATGCTGTTTTCTCCACCCAGTTAACCTCTCTATAACTGAATCTGCATGTGAAACACCTAGCTCGTTAGCACTGCAGGGGTTAAACGGGTTTGCCTGTTTCTCAGTACAGCTAGCACACTTACTGATACAAAGTGTCACATACGACTCATAAGTGGCAAACAGAATAATACTGGGGTTATTTTAAGGG

11. **TSS ΔSox17 *lhx5_*enhancer: Chr01:133702140-133702540**

>dSox17_lhx5

GGGTTACTGGCAGGATTGGTTTGGGCCGTATCTATACTTCACACAGGAGGGGGGTCTGGAATCGAACGCGCACCTAATCCAGATTTTGACTTCCTACAGGACAGTAGGAGGGAATTCCCTGTAGGACTCCCTAACCACCCCTATGTCGAAGGTAAATAAACTTAGCATTAAGCATGTGGCACCTACACTGCCCTTTCATAGAACTTCATTATGATGCTGTTTTCTCCACCCAGTTAACCTCTCTACCGGCATGTCTGCATGTGAAACACCTAGCTCGTTAGCACTGCAGGGGTTAAACGGGTTTGCCTGTTTCTCAGTACAGCTAGCACACTTACTGATACAAAGTGTCACATACGACTCATAAGTGGCAAGGCTAGCGCTACTGGGGTTATTTTAAGGG

12. **TSS ΔTcf *lhx5_*enhancer: Chr01:133702140-133702540**

>dTcf_lhx5

GGGTTACTGGCAGGATTGGTCAGAGTAACCGTGCAGAAAGAAACAGGAGGGGGGTCTGGAATTTTGTTTTCACCTAATCCAGATTTTGTATGTGAATAGGACAGTAGGAGGGAATTCCCTGTAGGACTCCCTAACCACCCCAAACACCAAGGTAAATAAACTTAGCATTAAGCAAACAATTAATACACTTACCATGACTAGAACTTCATTATGATGCTGTTTTCTCCACCCAGTTAACCTCTCTATAACTGAATCTGCATGTGAAACACCTAGCTCGTTAGCACTGCAGGGGTTAAACGGGTTTGCCTGTTTCTCAGTACAGCTAGCACACTTACTGATACAAAGTGTCACATACGACTCATAAGTGGCAAACAGAATAATACTGGGGTTATTTTAAGGG
